# Computational analysis to assess hemodynamic forces in descending thoracic aortic aneurysms

**DOI:** 10.1101/2024.11.22.624794

**Authors:** Francesca Duca, Daniele Bissacco, Luca Crugnola, Chiara Faitini, Maurizio Domanin, Francesco Migliavacca, Santi Trimarchi, Christian Vergara

**Affiliations:** LaBS, Dipartimento di Chimica, Materiali e Ingegneria Chimica, Politecnico di Milano, Milan, Italy; Department of Clinical Sciences and Community Health, Università degli Studi di Milano, Milan, Italy; Section of Vascular Surgery, Cardio Thoracic Vascular Department, Fondazione I.R.C.C.S. Ca’ Granda Ospedale Maggiore Policlinico, Milan, Italy; Mines Paris-PSL, Paris, France

**Keywords:** Thoracic aortic aneurysm, Fluid-Structure Interaction, Drag forces, TEVAR, Landing zones

## Abstract

Descending Thoracic Aortic Aneurysm (DTAA) is a life-threatening disorder, defined as a localized enlargement of the descending portion of the thoracic aorta. In this context, we develop a Fluid-Structure Interaction (FSI) computational framework, with the inclusion of a turbulence model and different material properties for the healthy and the aneurysmatic portions of the vessel, to study the hemodynamics and its relationship with DTAA.

We first provide an analysis on nine ideal scenarios, accounting for different aortic arch types and DTAA ubications, to study changes in blood pressure, flow patterns, turbulence, wall shear stress, drag forces and internal wall stresses. Our findings demonstrate that the hemodynamics in DTAA is profoundly disturbed, with the presence of flow recirculation, formation of vortices and transition to turbulence. In particular, configurations with a more steep aortic arch exhibit a more chaotic hemodynamics. We notice also an increase of pressure values for configurations with less steep aortic arch and of drag forces for configurations with distal DTAA.

Secondly, we replicate our analysis for three patient-specific cases (one for type of arch) obtaining conforting results in terms of accordance with the ideal scenarios. Finally, in a very preliminary way, we try to relate our findings to possible stent-graft migrations after TEVAR procedure to provide predictions on the post-operative state.

**KEY POINTS:**

- This study employs computational methods to assess hemodynamic forces in descending thoracic aortic aneurysms;
- We consider ideal cases by varying aortic arch type and aneurysm location;
- Our results show: chaotic hemodynamics for steep aortic arches; increase of pressure values for less steep aortic arches; high risk of plaque in the sac for proximal aneurysms and in the neck for distal aneurysms;
- We analyse also 3 patient-specific cases, confirming the major outcomes found for the ideal cases;
- We try to suggest how our pre-operative findings may be put in relation to assess the risk of stent-graft migration of a possible TEVAR procedure.

## 1 INTRODUCTION

Thoracic aortic aneurysm (TAA) is one of the most serious and potentially fatal diseases regarding the aorta. Descending Thoracic Aortic Aneurysm (DTAA) is defined as a localized increase in the descending segment of the Thoracic Aorta (TA) diameter of at least 50% with respect to the same aortic segment in age-matched and sex-matched healthy individuals (Bossone and Eagle, 2021). Even though DTAA pathophysiology is still poorly understood, from a histopathologic point of view, aneurysm formation may involve an alteration of the quantity and architecture of the tunica media components of Descending TA (DTA), namely elastin, collagen fibers and smooth muscle cells (Goldfinger et al., 2014). This degenerative process leads to a progressive remodeling, which induces a diminished aortic resilience and tensile strength, aortic wall thinning, increased wall stresses, and further aneurysm dilation, all of which could ultimately culminate in catastrophic events such as dissection and/or aneurysm rupture (Clift and Cervi, 2020).

Currently, there are two available surgical procedures for DTAA treatment: the open surgical strategy and the Thoracic Endovascular Aortic Repair (TEVAR). Specifically, TEVAR is a minimally invasive strategy based on aneurysm sac exclusion and depressurization through the deployment of a self-expandable stent-graft, and secure endograft fixation in correspondence of normal aorta proximal and distal zones of attachment, known as landing zones. This minimally invasive procedure represents a well-established alternative to open repair in individuals with suitable anatomic features, particularly in patients considered at high surgical risk but with a reasonable life expectancy after the procedure (Marrocco-Trischitta et al., 2009).

Clinical guidelines suggest surgical intervention for DTAA when its diameter is ≥ 60 mm or expands at a rate ≥ 5 mm/year (Riambau et al., 2017). However, there still remains a significant incidence of adverse events in patients whose aortas are smaller than these intervention thresholds (Elefteriades and Farkas, 2010). Thus, several prognostic factors have been discussed to better investigate and predict aneurysm progression and/or rupture (Fillinger et al., 2003; Venkatasubramaniam et al., 2004; Mourato et al., 2022). Among these, hemodynamic parameters comprise a useful tool in analyzing and quantifying the risk of negative outcomes in DTAA (Shang et al., 2013; Mandigers et al., 2023). In fact, aneurysm development has been associated with adverse vascular remodeling, induced by complex flow dynamics and abnormal Wall Shear Stress (WSS), which tends to be lower than in healthy aortic segments (Peattie et al., 2004). In particular, hemodynamics in DTAA is profoundly disturbed, presenting flow recirculation, vortex formation and adverse WSS, all of which possibly lead to further aneurysm growth and progressive aortic wall thinning, weakening, and even rupture (Tan et al., 2009b). Accordingly, the knowledge of the local hemodynamics in DTAAs may improve aneurysm expansion and rupture risk assessment.

Nowadays, computational modeling has become a valuable tool to simulate and analyse complex hemodynamics in the cardiovascular system, providing new insights into the mechanisms and pathophysiology of artery diseases, such as aortic aneurysm (Vorp et al., 1998; Boussel et al., 2008; Doyle et al., 2009; Boyd et al., 2016). Computational modeling can be used to precisely quantify the blood flow characteristics, wall forces and wall displacements in aortic aneurysms, either using a rigid wall assumption (Frauenfelder et al., 2006; Tse et al., 2011; Soudah et al., 2013) (Computational Fluid Dynamics - CFD) or a Fluid-Structure Interaction (FSI) approach (Di Martino et al., 2001; Scotti et al., 2005; Borghi et al., 2008).

Even though there are many works that focused on hemodynamics and WSS in presence of both abdominal (Wolters et al., 2005; Molony et al., 2009; Biasetti et al., 2010; Piccinelli et al., 2013) and ascending TAAs (Pasta et al., 2013; Callaghan et al., 2015; Campobasso et al., 2018; Etli et al., 2021), DTAAs have been poorly investigated by computational modeling. CFD experiments in Numata et al. (2016) showed for two DTAA patients very low velocity values within the aneurysm, as well as the presence of a vortical flow in systole, leading to transition to turbulence in the ascending aorta and high Oscillatory Shear Index (OSI) areas, which might be associated with atherosclerotic plaque formation. Duronio and Di Mascio (2023) employed CFD to compare the flow patterns in normal and DTAA subjects. Their findings demonstrated that in the aneurysmatic aorta blood flow was unstable with recirculation regions within the aneurysm, causing prolonged contact between blood particles and the aneurysm lumen surface, potentially leading to plaque deposition. When the vascular district undergoes relatively large displacements, like TA, FSI modeling may improve the hemodynamics description (Reymond et al., 2013), as well as provide information on the state of stress of the wall (Mendez et al., 2018) and on the disease progression (Valente et al., 2022). Tan et al. (2009b) demonstrated that FSI approach may provide a clearer insight into the development of DTAA, if compared with CFD modeling (Tan et al., 2009a), allowing to examine the effects of factors such as geometrical features and wall composition on blood velocity and wall stress. In this direction, Ong et al. (2019) compared hemodynamic quantities in a healthy and a DTAA case, whereas Silva et al. (2023) investigated the behaviour of the flow field and points of maximum stress and displacement of the arterial wall in aortic arch aneurysms. Dadras et al. (2023) exploited a FSI model to compare different hemodynamic and structural parameters before and after TEVAR, finding a significant improvement in velocity and pressure distribution and the damping of swirling flows strength after stenting.

Previous studies highlighted that the magnitude and orientation of drag forces may identify landing zones with a hostile biomechanical environment (Marrocco-Trischitta et al., 2019), which therefore may cause an insufficient proximal seal or migration of the endoprosthesis used in TEVAR (Marrocco-Trischitta et al., 2020). In this regard, while there are several studies that used CFD to investigate drag forces after the stent-graft deployment (Fung et al., 2008; Figueroa and Zarins, 2011; Krsmanovic et al., 2014), there are few works that focus on the pre-operative scenario (Marrocco-Trischitta et al., 2019; Belvroy et al., 2020), and none of these exploits FSI. Domanin et al. (2021) highlighted the predictive potential of the pre-operative analysis, describing a case report in which the proximal landing zone of the endoprosthesis was chosen according to the pre-operative computation of drag forces, by means of CFD, and six months follow-up CTA confirmed the validity of that decision.

In this study, we develop a FSI computational framework to investigate the flow patterns and the wall motion in twelve highly representative scenarios, encompassing the three type of aortic arch (i.e., Type I, Type II and Type III (Madhwal et al., 2008)) and three different aneurysmatic configurations, all obtained by virtual manipulation of Computed Tomography Angiography (CTA) images of one healthy patient. The first aim of this work consists in estimating, for each scenario, several hemodynamic quantities (i.e., blood pressure and velocity, turbulence, OSI and drag forces (Marrocco-Trischitta et al., 2018)), as well as vessel wall internal stresses. On the basis of this computational analysis, we are able to provide a comparison among the various scenarios, quantifying the differences between the pathological and the healthy cases, in terms of hemodynamic forces and wall stresses. Moreover, we also apply the same analysis to three patients with similar configurations to some of the idealized scanrios to demonstrate the validity of the results obtained in the latter cases. The second purpose of this work consists in exploiting the aforementioned pre-operative analysis with the aim of creating a link between pre-operative drag forces and the optimal landing zones for the stent-graft anchoring. Thus, for each diseased configuration, we are able to identify which landing zones might be the most hemodynamically disturbed and, for this reason, should be avoided for stent-graft sealing.

## 2 MATERIALS AND METHODS

### 2.1 Generation of computational scenarios

In this study, the pre-processing phase allows us to build *virtual ideal* and *patient-specific* geometries and meshes as described in what follows.

#### 2.1.1 Virtual geometries

Here we focus on the development of virtual thoracic aortic models with three distinct arch type (i.e., Type I, Type II and Type III), according to the Aortic Arch Classification (Madhwal et al., 2008), and differently located fusiform aneurysms along the descending portion of the vessel (i.e., proximal, central and distal). Thus, twelve Thoracic Aorta (TA) scenarios are developed: three healthy configurations, one for each arch type, and nine with a Descending Thoracic Aortic Aneurysm (DTAA) located in the proximal, central or distal Descending Thoracic Aorta (DTA), further divided according to the arch type.

The starting point is a healthy TA with a Type III aortic arch, obtained from the segmentation of Computed Tomography Angiography (CTA) images of a 40-years-old man taken at Fondazione IRCCS Ca’ Granda Ospedale Maggiore - Policlinico di Milano (all patient’s data are anonymized by a staff physician before the computational analysis). CTA scans were performed using a Siemens Somatom Definition Flash scanner (Siemens Healthineers, Forchheim, Germany) with the following acquisition parameters: slice thickness of 0.75 mm, reconstruction matrix of 512 × 512 pixel, and in-plane resolution of 0.396 mm × 0.396 mm. A level-set technique with a colliding fronts initialization, provided by the *Vascular Modeling Toolkit* (*vmtk*) (Izzo et al., 2018), is employed for the segmentation of the TA lumen from its ascending segment to the distal section, at the level of the celiac tripod, in order to comprise the first four thoracic *landing zones* of the Ishimaru’s map (Ishimaru, 2004). Then, by means of the *Paraview* (Ahrens et al., 2005) *filter Ruler*, the Left Common Carotid Artery (LCCA) diameter at its origin, and the distance between the on-set of the Brachiocephalic Artery (BCA) and the top of the arch are measured. Since the latter is more than two times greater than the diameter of the carotid, the aortic arch is classified as Type III according to the Aortic Arch Classification. The following step consists in virtually creating the other two arch types, namely Type I and II, varying the arch curvature, using *Autodesk Meshmixer* (Inc., 2019). *The main available functions adopted for this purpose are Transform*, for tilting in space the aortic arch, *Erase & Fill* and *Discard*, for progressively smoothing the modified surface; moreover *Remesh* tool is used to keep a high quality surface mesh, i.e., with fine triangulation, to avoid sharp edges while changing aortic curvature.

After checking that arch types had been reproduced properly, pathological aortic configurations are developed by exploiting *Boolean Union* of spheres in *Meshmixer*, and subsequently smoothing the bulging surface to create fusiform aneurysms with a maximum diameter of 60 mm, that is the critical value at which the incidence of aneurysm rupture, dissection, or patient death reaches the maximal level (Elefteriades and Botta, 2009). For each of the three arch models, the position of the virtually added aneurysmal dilation is varied distally along the descending portion of the vessel, thus leading to the creation of the nine DTAA cases.

Figure 1a shows the twelve configurations created following the aforementioned procedure. In what follows, these will be referred to as X-Y, where X = H, P, C, D indicates the location of the aneurysm (Healthy, Proximal, Central and Distal), whereas Y = I, II, III according to the arch type.

**FIGURE 1.**
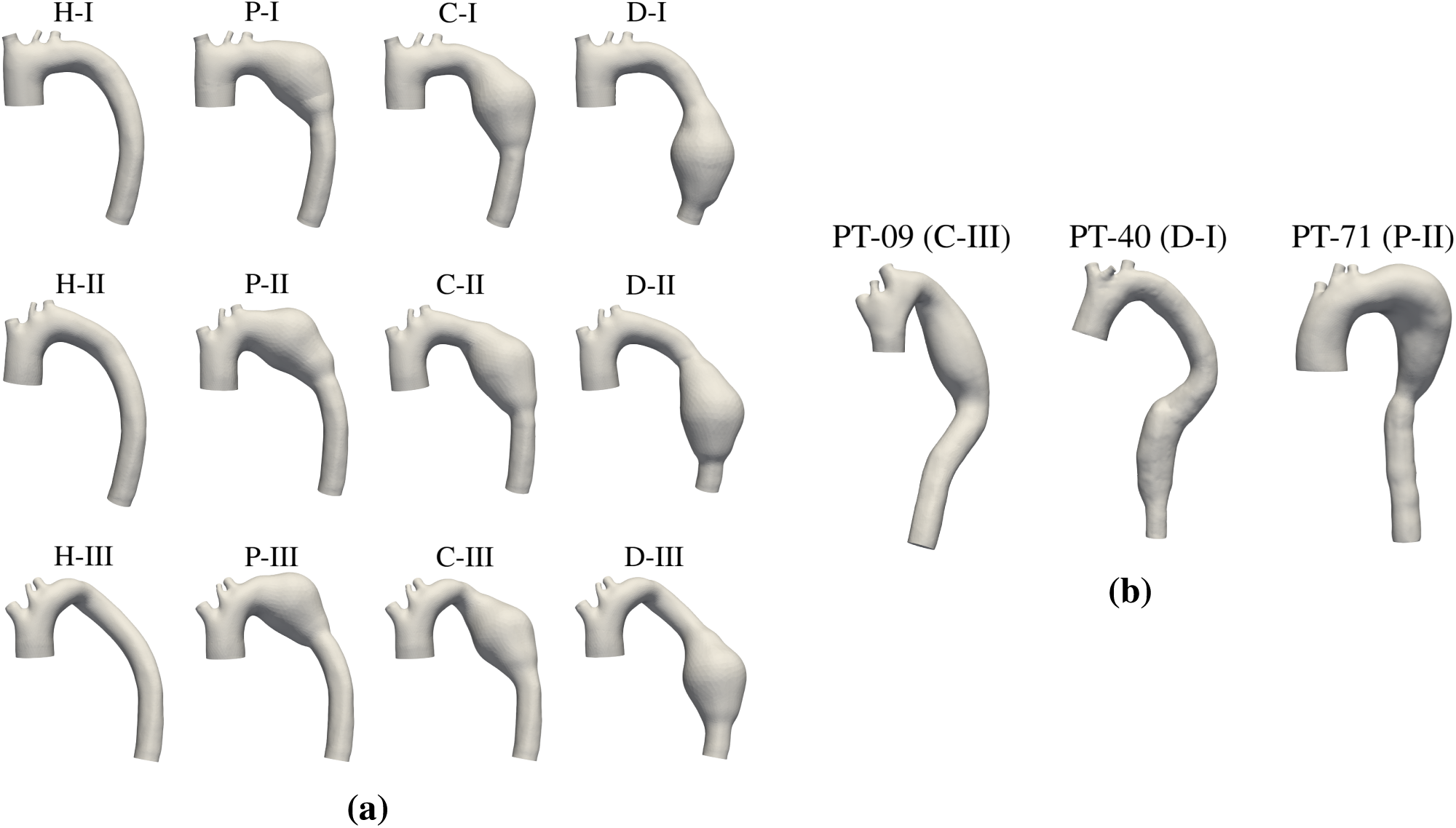
**(a)** Twelve virtual scenarios. Every virtual geometry is named by considering the location of aneurysm (H for healthy, P for proximal, C for central, D for distal) and arch type (I for Type I, II for Type II, III for Type III). **(b)** Three patient-specific configurations. PT-09: Type III with central aneurysm. PT-40: Type I with distal aneurysm. PT-71: Type II with proximal aneurysm. In the brackets we report the corresponding virtual scenario.

#### 2.1.2 Patient-specific geometries

Regarding the patient-specific configurations, we select three patients (PT-09, PT-40, PT-71) from Fondazione IRCCS Ca’ Granda Ospedale Maggiore - Policlinico di Milano dataset, who underwent pre-operative CTA scanning for TEVAR procedure planning. The acquisitions of CTA images were performed with a Siemens Somatom Definition Flash scanner (Siemens Healthineers, Forchheim, Germany) with the following acquisition parameters: slice thickness of 3.0 mm, reconstruction matrix of 512 × 512 pixels, and in-plane resolution of 0.629 mm × 0.629 mm. The patients are selected because they have a TA similar to some of the idealized configurations in terms of arch type and aortic tortuosity. In particular, PT-09, PT-40 and PT-71 present a central, distal and proximal DTAA, respectively. Vascular surgeons classified the aortic arch as Type III for PT-09, as Type I for PT-40, and as Type II for PT-71. The reconstruction of the patients’ TA lumen is performed by segmenting their pre-operative CTA scans using *vmtk* (as explained in Section 2.1). The three patient-specific geometries are shown in Figure 1b.

#### 2.1.3 Computational meshes

For all the twelve virtual surfaces and the three patient-specific configurations, the computational fluid meshes are generated exploiting *vmtk*. Specifically, the dimension of the meshes’ tetrahedral cells is defined as radius-dependent, so that the grid element size decreases as the diameter of the vessel diminishes, except for the aneurysm area, where a constant value equal to 1.5 mm is set. Moreover, close to the vessel wall, we consider a *Boundary Layer* (BL) composed by three sublayers with a total thickness of 0.5 mm for the supraortics and 1.0 mm for the rest of the domain.

Since the available CTA images do not allow us to detect the aortic wall, we create the meshes for the solid problem by extruding outward the fluid mesh’s external surface and selecting a wall thickness equal to 10% of the lumen radius for the healthy vessel, and equal to 1.5 mm (Van Puyvelde et al., 2016) for the aneurysmatic wall. In any case, a three-layered solid mesh is generated.

### 2.2 Mathematical modeling

In the present study, blood is assumed to be a Newtonian, homogeneous, and incompressible fluid, with a density of 1060 kg/m^3^ and viscosity of 3.5·10^−3^ Pa s; accordingly, its behavior is mathematically modeled with the Navier–Stokes equations written in the *Arbitrary Lagrangian–Eulerian* (ALE) formulation (Van de Vosse et al., 2003), where we use finite elasticity as lifting operator to recover the fluid domain configuration (Johnson and Tezduyar, 1994; Shamanskiy and Simeon, 2021). In case of pathological conditions, included DTAAs, transition to turbulence may be present and should be accounted for in the numerical simulations (Tan et al., 2009a). The σ-model *Large Eddy Simulation* (LES) turbulence model (Nicoud et al., 2011; Vergara et al., 2017) is used, since it is suitable for enclosed domains, such as vessels and ventricles.

Regarding the structure, the aortic wall is considered nearly incompressible and linearly elastic (Gao et al., 2006). The density of the aortic wall is set to 1000 kg/m^3^ while the Poisson’s ratio is set to 0.45. As the wall stiffness is a subject-specific parameter, a universal value does not exist. In this work, the Young’s modulus of the healthy part of the TA is set equal to 0.8 MPa, consistent with the literature (Lantz et al., 2011), while, since the aorta tends to stiffen when increasing its diameter (Vorp et al., 2003), the elastic modulus of the aneurysmatic wall is set equal to 1.2 MPa (Azadani et al., 2013). Thus, the aortic wall dynamics is formulated by means of the linear elasticity (Hooke’s law) written in the *Lagrangian configuration* (Donea et al., 2004), accounting for two different Young’s moduli for the healthy and the diseased parts.

Finally, the fluid and structure subproblems are coupled at their interface surface through: the *kinematic condition*, which provides the perfect adhesion between the fluid and structure particles; the *dynamic condition*, which prescribes the equilibrium between the fluid and structure forces; the *geometric condition*, which guarantees perfect adherence between the fluid and structure domains (Quarteroni et al., 2017).

Since each computational simulation is performed on a specific region of interest, which is a limited part of the whole cardiovascular system, the regions excluded from the model must be taken into account by exploiting suitable Boundary Conditions (BCs). Regarding the fluid problem, at the Ascending TA (ATA) inlet a physiological time-dependent pressure-wave ranging between 70 and 120 mmHg of 0.735 s period is imposed (Figure 2a) (Alastruey et al., 2012a). Moreover, in order to avoid the phenomenon of spurious reflections that may be caused by the artificial truncation of the computational domain, at each supraortics outlet (i.e., BCA, LCCA, and Left Subclavian Artery LSA), a proper absorbing boundary condition consisting of one resistance is prescribed (Nobile and Vergara, 2008) (Figure 2b on the left). Finally, at DTA outlet a 3-Element Windkessel Model (3-EWM) is imposed, since it provides a dynamic relationship between pressure and flow rate accounting for the downstream circulation (Westerhof et al., 2009) (Figure 2b on the left).

**FIGURE 2.**
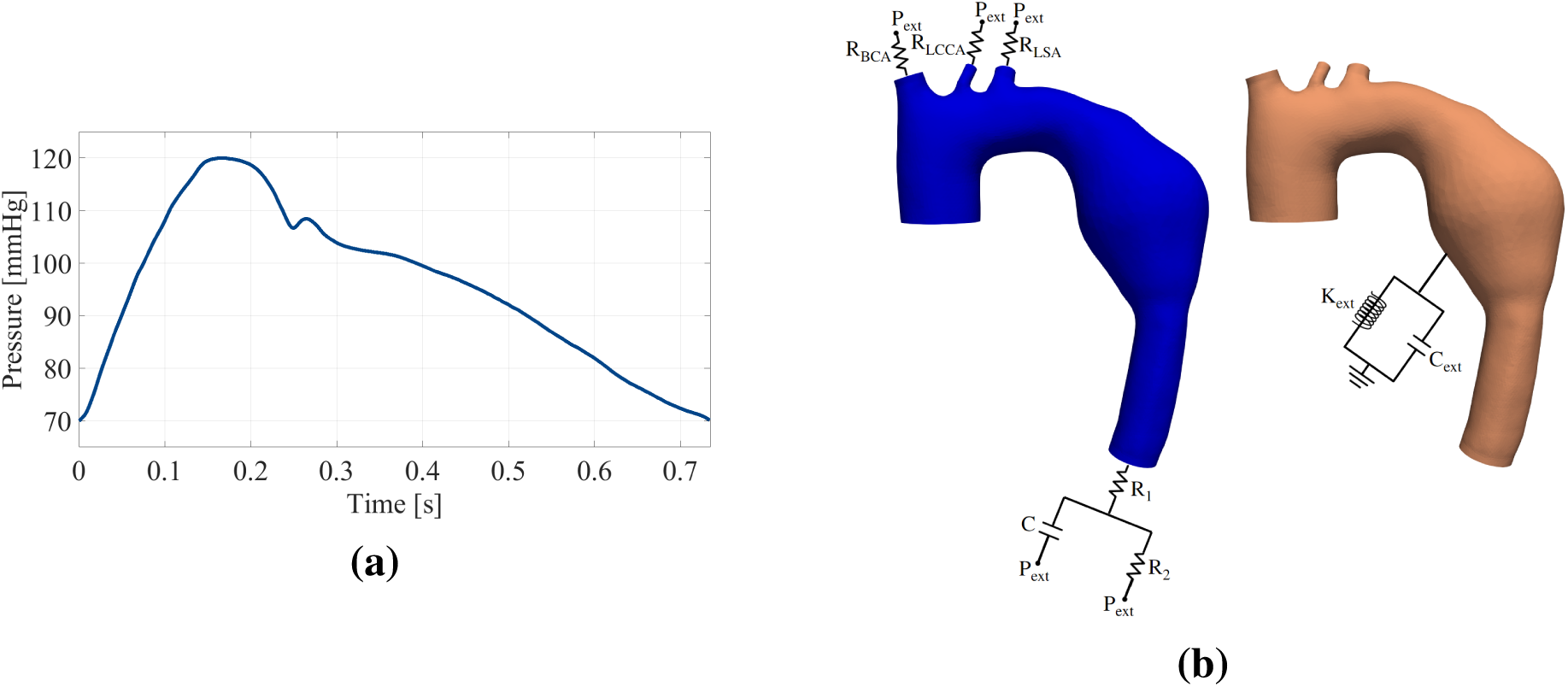
Boundary conditions prescribed for all the configurations (i.e., virtual and patient-specific). **(a)** Time-dependent inlet pressure waveform adapted from Alastruey et al. (2012a). **(b)** On the left: Absorbing and Windkessel boundary conditions prescribed at the outlets of the computational fluid domain. On the right: Robin boundary condition prescribed on the external surface of the computational structure domain.

Regarding the computational solid domain, at each inlet and outlet a null displacements condition is prescribed, whereas on the external surface, a Robin condition (Moireau et al., 2012) with springs of stiffness K_ext_ is supplied by a dissipative element of capacity C_ext_ (Regazzoni et al., 2022) (Figure 2b on the right) modeling the constraint exerted by the surrounding tissue (e.g., the spinal column) on the vessel’s movements.

### 2.3 Numerical approximation and setting of the numerical simulations

Regarding the numerical approximation of the Fluid-Structure Interaction (FSI) problem, for the time discretization we impose a time-step Δt = 5·10^−4^ s, and we consider the semi-implicit Backward Differentiation Formula of second-order (BDF2) for the fluid subproblem, whereas first-order BDF1 for the structure problem. At time *t*^*n*+1^, in the σ-LES model, we compute the turbulent viscosity using the 2-order extrapolation of fluid velocity from the previous time steps (Nicoud et al., 2011). Moreover, both the absorbing and 3-EWM BCs are coupled explicitly to the Navier-Stokes equations, i.e., the boundary pressure at time *t*^*n*+1^ is computed exploiting the flow rate calculated from the previous time step *t*^*n*^ (Pozzi et al., 2021). At each time step *t*^*n*+1^, we solve first the lifting problem with boundary displacement obtained from the structure problem at time *t*^*n*^; then, once the new fluid domain is computed, the FSI problem is solved monolithically by using GMRES preconditioned by means of a block preconditioner (see Bucelli et al. (2023)).

For each configuration reported in Figure 1, the simulation is performed over ten cardiac cycles of which we discard the first three to allow the stabilization of flow velocity and pressure fields. Indeed, after four cycles, the numerical results reach a regime configuration with, however, fluctuations among heartbeats (HBs) due to transition to turbulence.

All the numerical simulations are performed using life^x^ (Africa, 2022; Africa et al., 2024): a high-performance library for the finite element simulations of multiphysics, multiscale, and multidomain problems, developed at the MOX, Dipartimento di Matematica, with the collaboration of LaBS, Dipartimento di Chimica, Materiali e Ingegneria Chimica, both at Politecnico di Milano.

As preliminary step before running the final simulations, a calibration of the BCs parameters is necessary. In particular, for all the parameters, a literature formula is used to propose an initial guess. Specifically, for the supraortics, the absorbing resistances R_BCA_, R_LCCA_ and R_LSA_ are computed according to geometric and mechanical quantities, using the formula reported by Nobile et al. (2013):

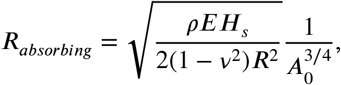

where *ρ* is the blood density, E is the Young’s modulus of the TA, *H*_s_ is the aortic wall thickness, ν is the Poisson ratio, *R* is the outlet radius, and *A*_0_ is the outlet area in the undeformed configuration. For the calibration of the 3-EWM parameters, the proximal resistance *R*_1_ is set as the one obtained in the case of a resistance absorbing boundary condition; the distal resistance *R*_2_ is set according to Romarowski et al. (2018), basically subtracting *R*_1_ to a total peripheral resistance, computed as the ratio between the time-averaged inlet pressure and flow rate:

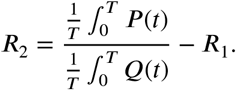

Instead, the compliance C is computed following the formula reported by Alastruey et al. (2012b):

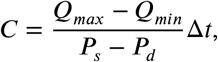

where *Q*_min_ and *Q*_max_ are the minimum and maximum blood flow rates in a cardiac cycle at the inlet, *P*_s_ and *P*_d_ are the systolic and diastolic pressure (all these values are taken from Alastruey et al. (2012a)) and Δt is the distance in time between *Q*_min_ and *Q*_max_. The downstream pressure P_ext_ (see Figure 2b on the left) is set to 70 mmHg in accordance with the values estimated by Alastruey et al. (2016). The initial guesses obtained through the formulas reported above are subsequently tuned, performing several simulations using the H-I configuration, in order to ensure that approximately 75% of the cardiac output entering in ATA exits through DTA (Coats, 1990). The two Robin parameters in the external structure BC are guessed starting from the values reported in Moireau et al. (2012), and then calibrated in order to guarantee a physiological cross-sectional area variation of about 10% during a cardiac cycle (Van Prehn et al., 2009). Table 1 shows the final calibrated values.

**TABLE 1.** Values of the calibrated parameters imposed in the outlet boundary conditions for the supraortic branches (BCA, LCCA, LSA) and for DTA.

|  |  |
| --- | --- |
| $R_{BCA}$ [kg/(s m<sup>4</sup>)] | $1.50 \cdot 10^8$ |
| $R_{LCCA}$ [kg/(s m<sup>4</sup>)] | $7.50 \cdot 10^8$ |
| $R_{LSA}$ [kg/(s m<sup>4</sup>)] | $5.50 \cdot 10^8$ |
| $R_1$ [kg/(s m<sup>4</sup>)] | $2.30 \cdot 10^7$ |
| $R_2$ [kg/(s m<sup>4</sup>)] | $6.08 \cdot 10^7$ |
| $C$ [(m<sup>4</sup> s<sup>2</sup>)/kg] | $8.00 \cdot 10^{-7}$ |
| $K_{ext}$ [Pa/m] | $5.50 \cdot 10^6$ |
| $C_{ext}$ [(Pa s)/m] | $4 \cdot 10^5$ |

### 2.4 Mesh sensitivity analysis

To allow reliable elaboration of Wall Shear Stress (WSS) values obtained by the numerical simulations, the blood flow close to the wall has to be accurately captured and hence, the first layer of the fluid mesh grid must be sufficiently close to the wall. This distance from the wall is represented as a nondimensionless distance *y*^+^ which, according to the turbulent BL sublayer coordinates (Wilcox et al., 1998), should be less than 2 in order to guarantee the correct WSS detection. Therefore, once we have chosen the thickness of the first BL sublayer *T*_1_, we compute *y*^+^ exploiting the formula (Febina et al., 2018):

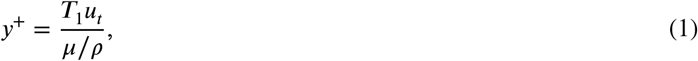

where 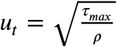 is the friction velocity, in which *τ*_max_ is the maximum WSS value during a cardiac cycle. Since the total BL thickness is diffe^*ρ*^rent for TA and the supraortics (see Section 2.1.3), and consequently also the thickness of the first sublayer, we ensure that *y*^+^ is less than 2 for both parts of the geometry (Table 2).

**TABLE 2.** Boundary layer sensitivity analysis results.

| $T_{aorta}^1$<br>[mm] | $\tau_{max}^{aorta}$<br>[Pa] | $y_{aorta}^+$<br>[-] | $T_1^{sup.}$<br>[mm] | $\tau_{max}^{sup.}$<br>[Pa] | $y_{sup.}^+$<br>[-] |
| --- | --- | --- | --- | --- | --- |
| 0.143 | 13.19 | 1.11 | 0.071 | 17.13 | 0.44 |

Subsequently, a sequence of three finite element meshes of increasing density is used to perform a mesh sensitivity analysis, in order to establish the most suitable fluid and solid mesh grid size, which ensures the correct balance between accuracy of the results and runtime of the simulations, and to avoid grid resolution errors. Thus, each simulation is run on three different couple H-I fluid and solid meshes (i.e., M1, M2 and M3) with a grid size *h*_fluid_ and *h*_solid_ progressively finer. The simulations are run for two HBs with the same inlet pressure-wave (Figure 2a) and only the results of the second cycle are considered. The results are declared mesh independent when the DTA outlet systolic peak flow rate (*Q*_peak_), peak WSS (*τ*_peak_) and peak wall displacement (*d*_peak_) between successive meshes satisfy the following relation:

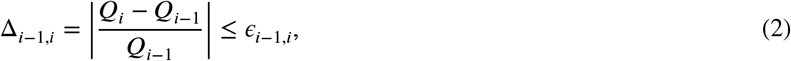

where *Q* is the quantity of interest evaluated for the coarser (*i*−1)-th and the finer *i*-th mesh, and *ϵ*_*i*−1,*i*_ is the tolerance computed as follows:

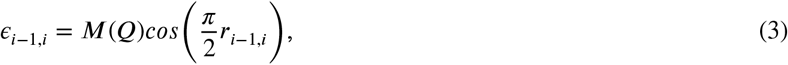

where 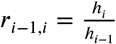 is the refinement ratio between the coarser (*i*−1)-th and the finer *i*-th mesh size, and *M*(*Q*) is an acceptable clinical error, p^*h*^r^*i*−1^escribed according to the accuracy associated with a specific diagnostic technique (e.g., echo color Doppler accuracy in measuring blood flow velocity). Table 3 summarizes the size, the number of cells and the quantities of interest for all the meshes considered.

**TABLE 3.** Grid size, number of cells and computed quantities of interest among the three meshes. *h*_fluid_: average size of the fluid mesh grid. *h*_solid_: average size of the solid mesh grid. *N*_fluid_: number of cells of the fluid mesh. *N*_solid_: number of cells of the solid mesh. *Q*_peak_: DTA outlet blood flow rate at the systolic peak. *τ*_peak_: WSS at the systolic peak. *d*_peak_: wall displacement at the systolic peak.

| Mesh | $h_{\text{fluid}}$<br>[mm] | $h_{\text{solid}}$<br>[mm] | $N_{\text{fluid}}$<br>[-] | $N_{\text{solid}}$<br>[-] | $Q_{\text{peak}}$<br>[ml/s] | $\tau_{\text{peak}}$<br>[Pa] | $d_{\text{peak}}$<br>[mm] |
| --- | --- | --- | --- | --- | --- | --- | --- |
| M1 | 1.568 | 1.573 | 957 634 | 283 431 | 271.82 | 11.85 | 6.10 |
| M2 | 1.298 | 1.302 | 1 307 670 | 336 568 | 273.25 | 13.19 | 6.11 |
| M3 | 1.036 | 1.040 | 2 004 246 | 553 546 | 273.51 | 13.60 | 6.15 |

Table 4 shows the results of the mesh sensitivity analysis, reporting the relative difference of a certain quantity *Q* between two consecutive meshes and the prescribed tolerances. Cells colored in green show that the relative difference of a certain *Q* is less than the tolerance, vice versa cells colored in red show that the difference is greater than the tolerance. Accordingly, the Table shows that the most suitable choice is M2, as its grid size is sufficiently fine to ensure that the quantities differences between M2 and M3 are below the imposed tolerance for both fluid and solid meshes, while it almost halves the computational time. In fact, whilst the total time required to simulate one HB with M1 is 4 hours, M2 needs approximately 6 hours against 14 hours required for M3.

**TABLE 4.**
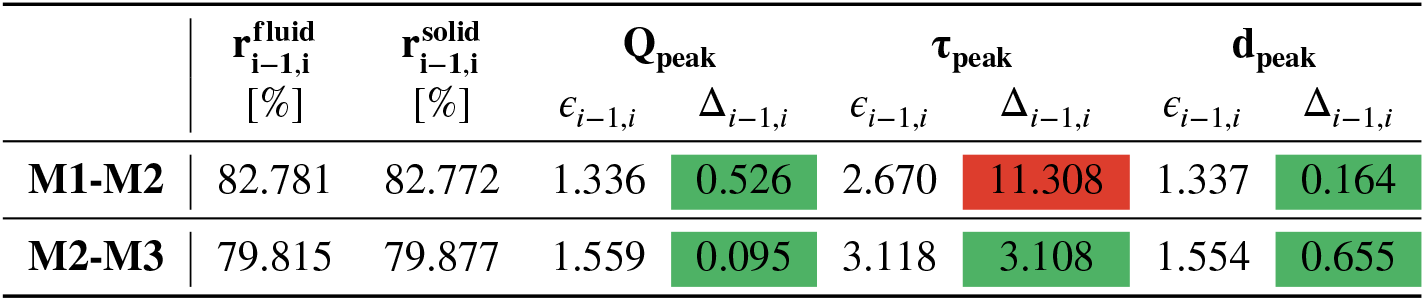
Mesh sensitivity analysis based on the computed quantity of interest, i.e., DTA systolic peak blood flow rate (*Q*_peak_), peak WSS (*τ*_peak_) and peak wall displacement 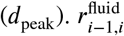 and 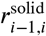 denote the refinement ratio between the coarser (*i*−1)-th and the finer *i*-th mesh. The specific tolerances *ϵ*_*i*−1,*i*_ and the relative difference of each quantity Δ _*i*−1,*i*_ between two consecutive meshes are computed following Equation (3) and Equation (2), respectively. The acceptable error *M* (*Q*) is set to 5% for *Q*_peak_ (Powell et al., 2000) and *d*_peak_, while is set to 10% for *τ*_peak_ (Pantos et al., 2007). All the reported quantities are percentages and *i* = 1, 2, 3.

| | $r_{i-1,i}^{\text{fluid}}$<br>[%] | $r_{i-1,i}^{\text{solid}}$<br>[%] | $Q_{\text{peak}}$ | | $\tau_{\text{peak}}$ | | $d_{\text{peak}}$ | |
| --- | --- | --- | --- | --- | --- | --- | --- | --- |
| | | | $\epsilon_{i-1,i}$ | $\Delta_{i-1,i}$ | $\epsilon_{i-1,i}$ | $\Delta_{i-1,i}$ | $\epsilon_{i-1,i}$ | $\Delta_{i-1,i}$ |
| **M1-M2** | 82.781 | 82.772 | 1.336 | 0.526 | 2.670 | 11.308 | 1.337 | 0.164 |
| **M2-M3** | 79.815 | 79.877 | 1.559 | 0.095 | 3.118 | 3.108 | 1.554 | 0.655 |

### 2.5 Quantities of interest

#### 2.5.1 Pre-operative analysis

To comprehensively observe and describe the hemodynamics and wall motion in all the configurations, we quantitatively evaluate the blood pressure and velocity, and structural displacement computed by the numerical simulations. Specifically, we compute the *ensemble* values of such quantities, i.e. the average calculated, at each time and at each point of the domain, over the seven HBs. Moreover, we estimate the following quantities:

- Standard Deviation (SD) at each time and space of the blood velocity with respect to its ensemble value. This allow us to quantify and localize the regions with considerable large velocity fluctuations, which are characterised by marked transition to turbulence Vergara et al. (2017);
- (Ensemble) Oscillatory Shear Index (OSI): a dimensionless descriptor which allows the identification of zones of the artery wall subjected to large directional variations of the WSS vector during a cardiac cycle, computed as follows (Ku et al., 1985):

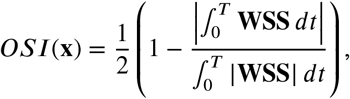

where **WSS** is the (ensemble) WSS vector and *T* is the cardiac cycle duration. In aneurysms, high values of OSI favour the development of atherosclerosis, inflammation of the artery and internal thickening of the arterial wall (Mutlu et al., 2023);
- (Ensemble) Von Mises Stress (VMS): a structural quantity that allows for an evaluation of the stresses a sample material is subjected to, in order to establish the response of the former to a certain complex loading condition. VMS is commonly used as a criterion for material failure and to predict the effect of the hemodynamics on the internal wall of aneurysms, associating high value of VMS to the possibility of aneurysm rupture Tan et al. (2009b). Exploiting the assumption of linear elasticity introduced for the aortic mechanics modeling (see Section 2.2), we are able to compute VMS, σ_*V M*_, with the formula:

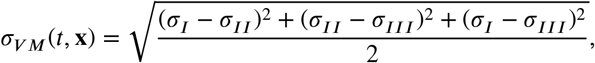

where σ_*i*_, with *i* = *I, II, III*, are the principal stresses of the Cauchy stress tensor in the i-th direction related to the Hook’s law.
- (Ensemble) *drag forces*: these are the forces (i.e., pressure and WSS) exerted by the blood on the vessel wall. For each of the geometries considered in this study (see Figure 1) we consider a subdivision, coming from the clinical practice, useful to assess how the state of stress along the vessel is distributed. Specifically, TA is subdivided into landing zones according to the Modified Arch Landing Areas Nomenclature (MALAN) classification Marrocco-Trischitta et al. (2017) (see Figure 3a), which merges Ishimaru’s map (Ishimaru, 2004) with the Aortic Arch Classification (Madhwal et al., 2008). We are here interested in considering the (vectorial) average of drag forces onto each landing zone surface *LZ* (refer to Figure 3b) (Marrocco-Trischitta et al., 2018):

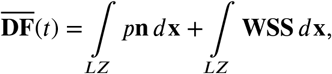

where **n** is the outward normal evaluated at the wall. Notice that, 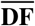 is a variable which, at each time t, assumes different constant values over the different landing zones. With 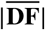 we will indicate its magnitude;
- (Ensemble) *DF*_∥_ and (ensemble) *DF*_⊥_: parallel (tangential) and normal (perpendicular) main components of 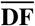 normalized over the area *A* of each landing zone:

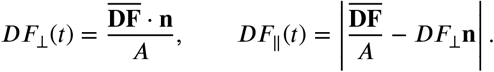

*DF*_∥_ is aligned with the aortic centerline, while *DF*_⊥_ acts orthogonally to the wall of the vessel, in the outward direction (Figure 3b).

**FIGURE 3.**
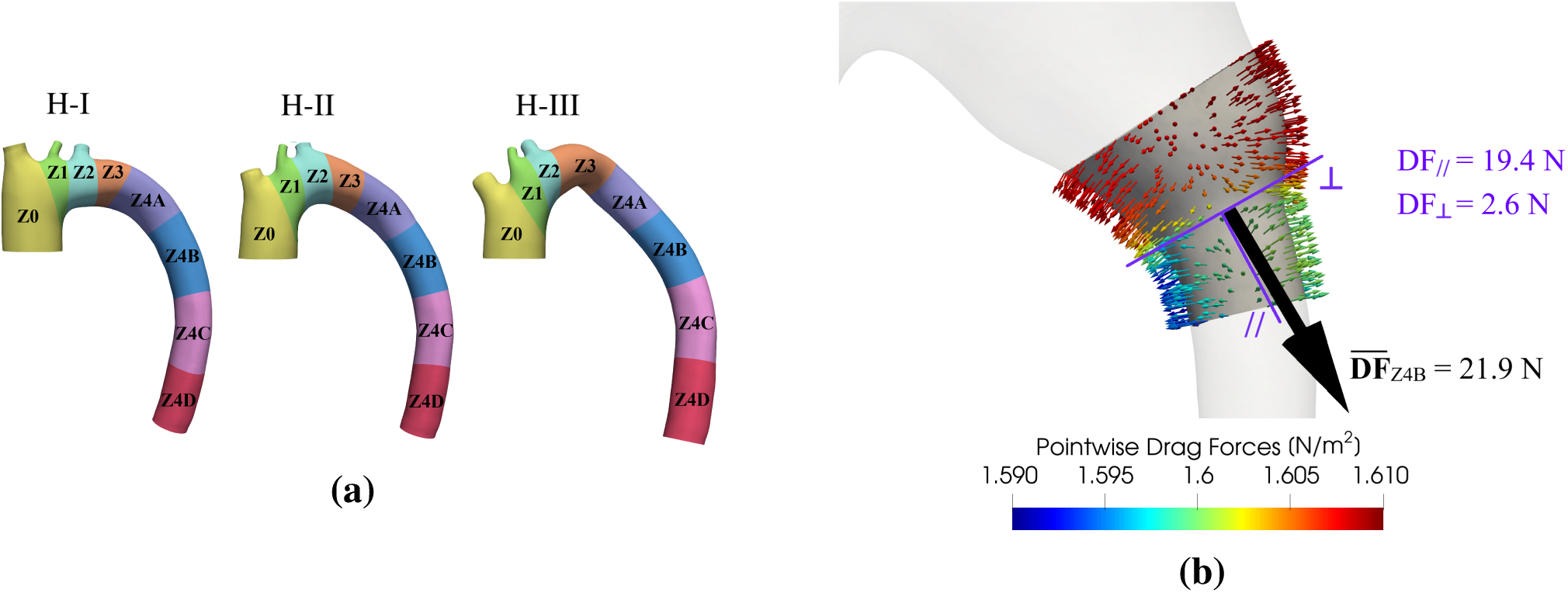
**(a)** Landing zones map for each aortic arch type according to the MALAN classification. **(b)** Example of the evaluation of *DF*_∥_ and *DF*_⊥_ in landing zone Z4B for P-II configuration. Coloured arrows: pointwise (ensemble) drag forces. Black arrow: average drag force 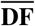 at the systolic peak. The direction ∥ tangent to the centerline and its outward perpendicular direction ⊥, used to determine the *DF*_∥_ and *DF*_⊥_ components, respectively, are shown in purple.

#### 2.5.2 Towards a preliminary TEVAR analysis

In what follows, we want to propose an index in view of the possible application of our pre-operative analysis to the case of Thoracic Endovascular Aortic Repair (TEVAR) procedure. The introduction of *DF*_∥_ and *DF*_⊥_ aims also at providing additional useful information to clinicians who are interested in predicting unwanted TEVAR late complications, especially the endograft migration. In view of preventing the latter risk, we discuss four possible mechanisms:

i. it would be better to avoid landing zones in which *DF*_∥_ results to be noticeably high, as it could promote the migration of the stent-graft, which is subjected to a significant force in the direction of the flow (∼ *DF*_∥_);
ii. it would be better to avoid the landing zones where *DF*_∥_ (independently of its actual value) results to be predominant over *DF*_⊥_ (Domanin et al., 2021) and, for this reason, we consider important the ratio *DF*_∥_/*DF*_⊥_ since, for the same *DF*_∥_, the condition whith larger *DF*_⊥_ is to be preferred, as it ensures that the stent-graft would be subjected to a sufficient radial force to gurantee its safe anchoring (∼ *DF*_∥_/*DF*_⊥_);
iii. we speculate that too high values of *DF*_⊥_ could be dangerous, as they may cause an excessive vessel dilatation, leading to the stent-graft detachment and, at the level of the aortic arch, to *endoleak* formation, particularly the so-called *bird-beak configuration* (Auricchio et al., 2014), *which may be a further source of stent-graft migration (Thomas and Sanchez, 2009)* 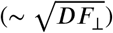;
iv. *we consider as unfavourable the situation in which the presence of the aneurysm causes a considerable increase of the parallel component with respect to the healthy condition, used as reference for a normal scenario* 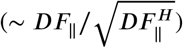

In order to take into account all the dangerous conditions mentioned above and to provide valuable insights about which landing zones would be the safest in view of the stent-graft anchoring, we propose the following sinthetic index *β (risk factor*), which is computed for each landing zone as follows:

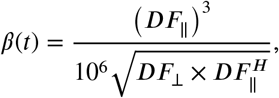

where 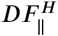 is the parallel component of the average drag forces in the healthy configuration.

## 3 RESULTS: VIRTUAL CONFIGURATIONS

In this Section, we report the results of the numerical simulations performed in all the virtual configurations. First, we report the results in terms of average quantities for the healthy and for one representative DTAA ideal case (Section 3.1); then, we report the 3D results of the simulations conducted in the virtual cases (Section 3.2), with a focus on drag forces (Section 3.3).

### 3.1 Average quantities

In Figure 4 we report the average blood flow rate and the cross-sectional area variation within one HB for H-I and C-I scenarios, selected here as representative for the DTAA cases.

**FIGURE 4.**
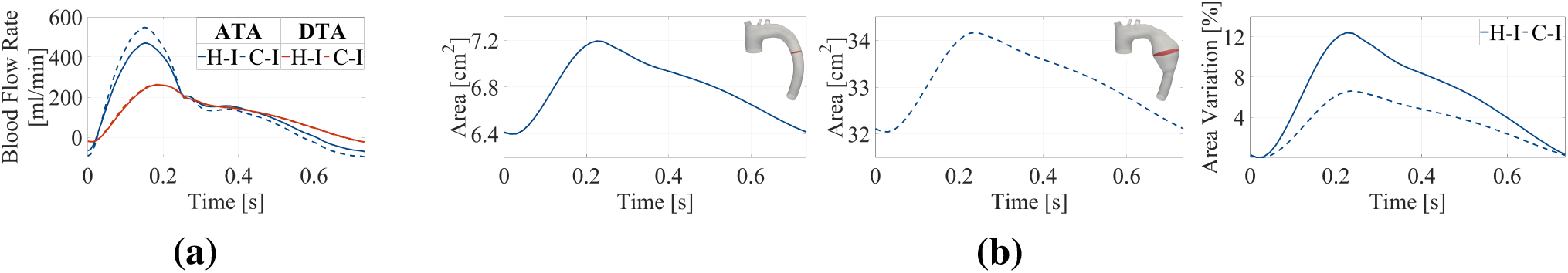
**(a)** Blood flow rates at the inlet of ATA and at the outlet of DTA during one cardiac cycle for H-I and C-I configurations. **(b)** From left to right: cross-sectional area over time in H-I configuration; cross-sectional area over time in C-I configuration; percentual variation of cross-sectional area in both H-I and C-I configurations. The boxes on the top-right corner highlight in red the cross-section considered.

First, we notice that our simulations are able to satisfy for the healthy scenario the two criteria used for the outflow BCs calibration (75% of inflow reaches the DTA outlet and area variation in time of about 10%, see Section 2.3). Indeed, we have a plausible average-in-time inlet blood flow rate of 6.6 L/min (Critchley and Critchley, 1999), against 5.1 L/min at the DTA outlet, corresponding to 77% of the inflow, while the remaining 23% goes to the neck vessels, in accordance with literature data (Coats, 1990). This corresponds to the plots in time reported in continuous lines in Figure 4a. The negative flow at the end of the cycle (about 6% of the ATA flow rate) is plausibly the blood flowing to the coronary arteries supplying the myocardium. As regards the second criterion, we notice from Figure 4b that, for the H-I configuration, the maximum area variation is 12.4%, in good accordance with the literature values (Van Prehn et al., 2009).

Secondly, in what follows we discuss the differences obtained between the results of H-I and C-I scenarios. Regarding the blood flow rate, in the diseased scenario, we have the same average-in-time value than in the healthy case (6.6 L/min), as a consequence of the same inlet pressure and outlet resistances and compliance in the two cases. However, from Figure 4a we notice that, in C-I scenario, the flow rate at the systolic peak is higher (17% more) than in the healthy configuration. Instead, for the area variation, we notice percentage values that are almost half (6.6%) of the healthy scenario, in accordance with the increased Young’s modulus in the aneurysm of C-I configuration.

### 3.2 Comparison of hemodynamics and structural stresses

In what follows, we report 3D maps of the ensemble quantities of interest obtained in the numerical simulations, such as pressure and velocity fields, velocity SD for turbulence quantification, OSI, and VMSs, with the aim of comparing the different hemodinamic answers of the virtual DTAA scenarios described in the previous Section.

#### 3.2.1 Pressure and velocity fields

Figure 5a shows the ensemble blood pressure over seven HBs at the systolic peak for all the virtual configurations. We observe that, while H-I and H-II scenarios have nearly the same trend, in H-III a region of relative low pressure is clearly visible immediately after the supraortics, where the curvature of DTA is maximum (highlighted by the black box in Figure 5a). Regarding the diseased scenarios, the pressure within the whole aneurysm is uniformly higher than the values found in the other parts of the geometry, and there is also a sudden rise and decrease in pressure values at the proximal and distal neck, respectively (Figure 5b). These findings are in good agreement with what reported by Campobasso et al. (2018), who proved that an increase in the aortic stiffness could cause an intensification of pressure values in ascending TAAs. The blood pressure is particularly high in C-I, C-II, D-I and D-II configurations, in contrast to the other scenarios, e.g. C-III and D-III, where again low pressure values are observed in the region of maximum DTA curvature.

**FIGURE 5.**
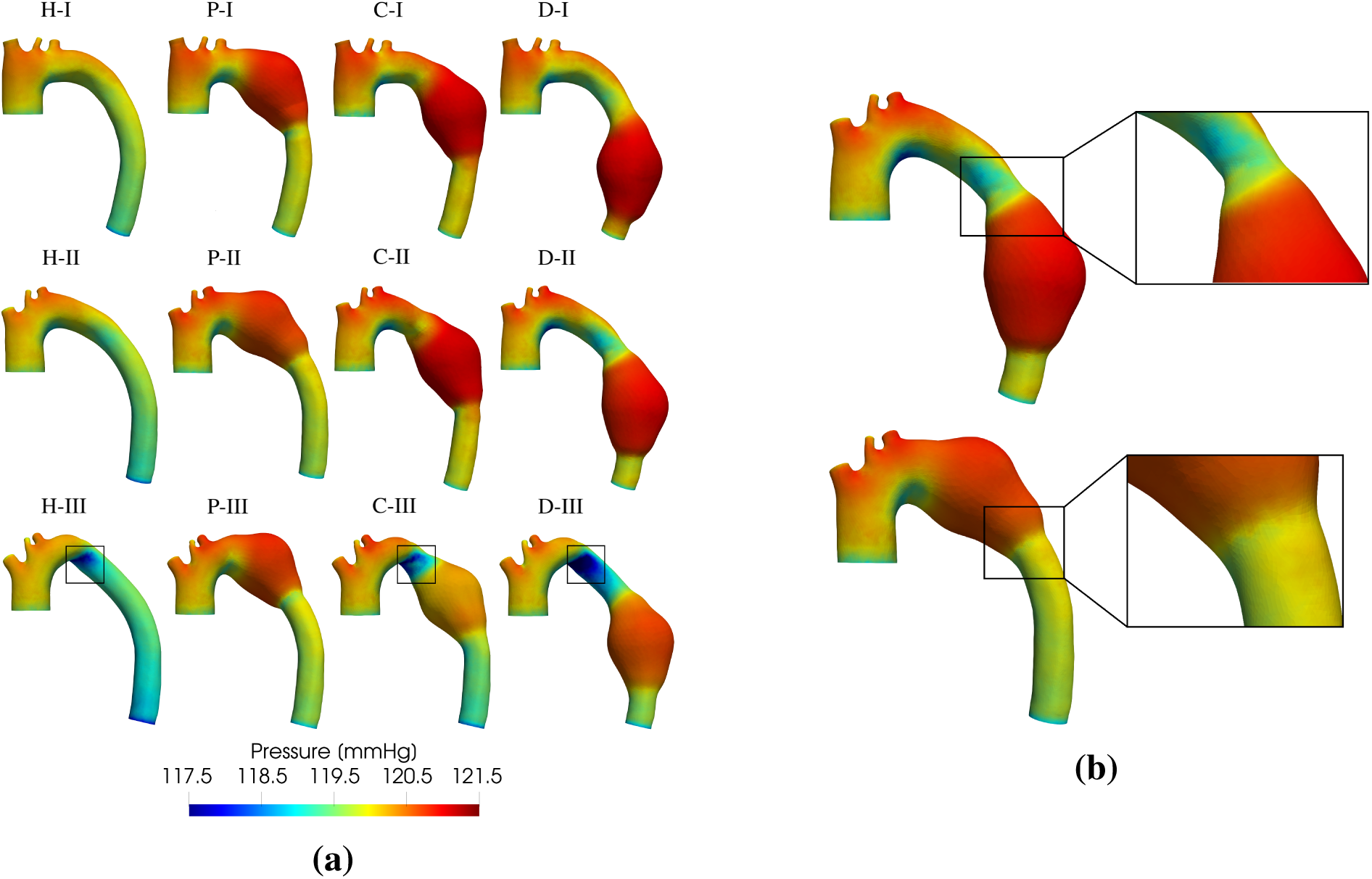
**(a)** Ensemble blood pressure at the systolic peak (t = 0.165 s) in all the virtual configurations. Black bloxes highlight the low pressure region in the inner curvature of the arch of the Type III configurations. **(b)** Top: sudden increase of pressure value at the proximal aneurysm neck in D-II configuration (chosen as a representative diseased case); bottom: sudden decrease of pressure value at the distal aneurysm neck in P-II configuration (chosen as a representative diseased case).

In Figure 6a we show the ensemble velocity magnitude over seven HBs at the systolic peak for all the configurations considered. It is possible to notice that, in the healthy tracts, the blood flow is very stable, characterised by smooth and almost parallel streamlines. On the contrary, the presence of the aneurysmal sac causes the flow to be slightly more chaotic, with recirculation regions at the proximal aneurysm neck in all the cases and with another vortical structure, adjacent to the inner side wall within the aneurysm, in C-III (black box in Figure 6a). Besides, while the ensemble velocity magnitude considerably decreases within the aneurysm, it increases in the regions just upstream and downstream the bulging if compared to the healthy cases, particularly in D-II, C-III and D-III configurations. In particular, in the region not directly affected by the pathology blood velocity records values up to 0.70 m/s, whereas in the aneurysmal sac it decreases greatly, reaching values between 0.10 and 0.30 m/s, in good accordance with what found by Silva et al. (2023).

**FIGURE 6.**
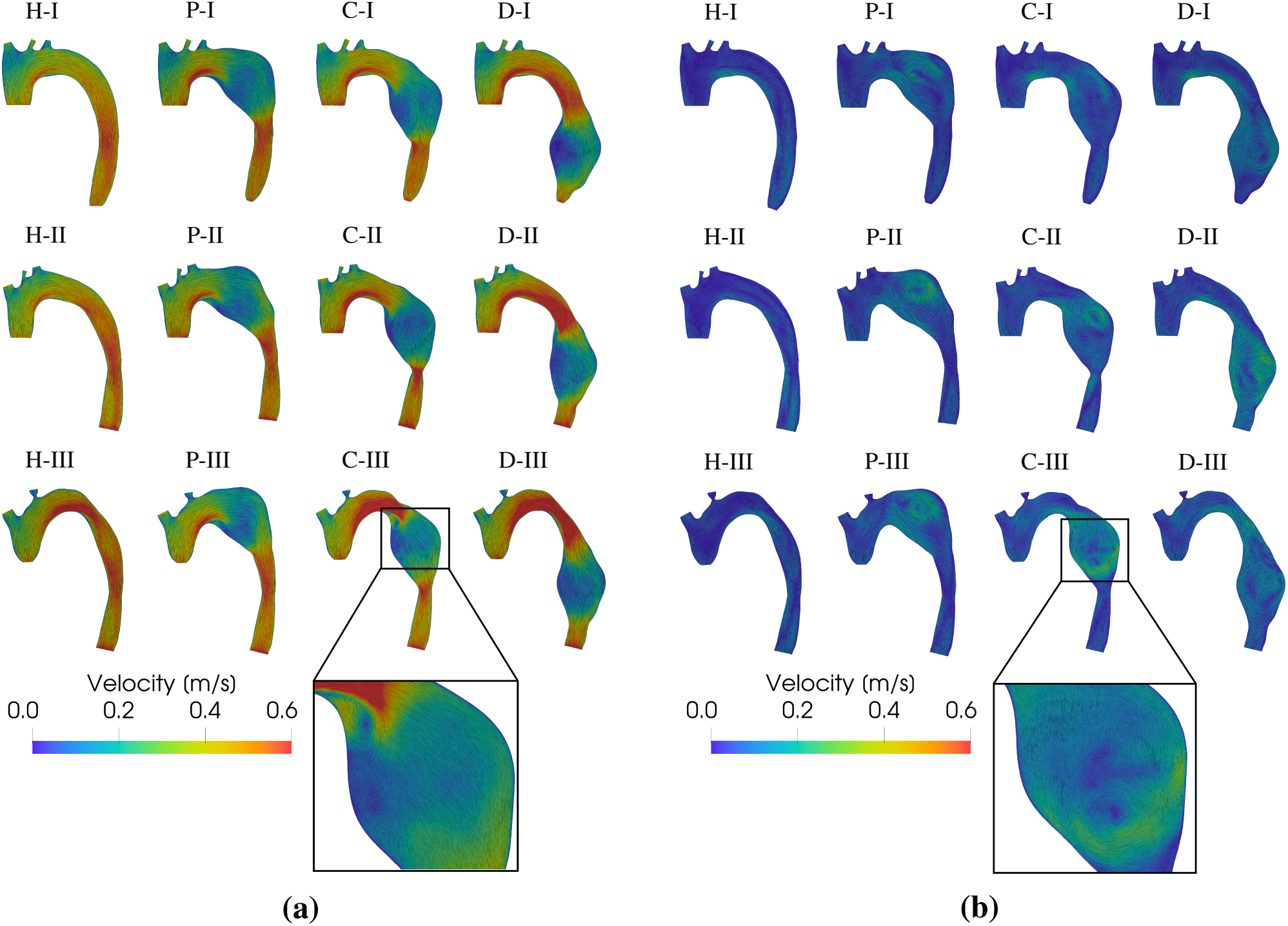
**(a)** Ensemble velocity magnitude over seven HBs at the systolic peak (t = 0.165 s) in all the twelve virtual configurations. Black box highlights the two vortices forming at the proximal aneurysm neck and on the inner side wall within the aneurysm in C-III configuration. **(b)** Ensemble velocity magnitude over seven HBs in late diastole (t = 0.645 s) in all the twelve virtual configurations. Black box highlights the vortex forming within the aneurysm in C-III configuration (chosen here as representative).

In Figure 6b we consider the diastolic phase, observing that in the aneurysm, blood velocity remains higher than in the healthy cases with large vortices (black box in Figure 6b), which are more evident in P-II, C-II, D-II, P-III and C-III configurations. The vortical structures forming within the aneurysm show velocity values ranging from 0.30 and 0.40 m/s, consistent with what reported by Silva et al. (2023).

#### 3.2.2 Transition to turbulence

In Figure 7 we report the SD of the velocity magnitude over seven HBs for all the virtual configurations considering three time instants: early systole (Figure 7a), late systole (Figure 7b) and late diastole (Figure 7c). Large SD values in a region represent high velocity fluctuations among HBs in that region. This is an index of transition to turbulence (Vergara et al., 2017).

**FIGURE 7.**
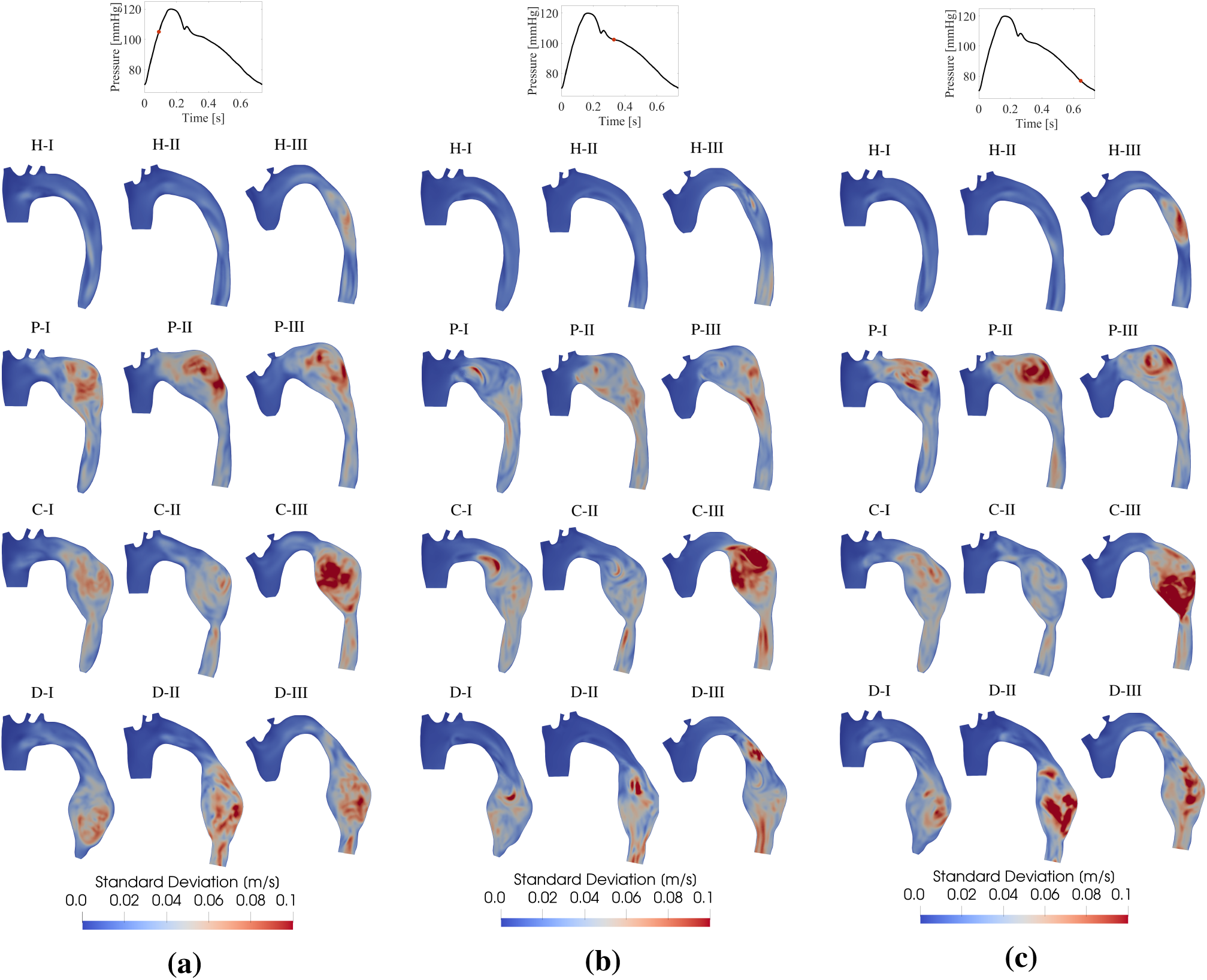
Standard Deviation (SD) of the velocity magnitude over seven HBs for all the virtual configurations at three time instants: **(a)** early systole t = 0.09 s, **(b)** late systole t = 0.33 s, **(c)** late diastole t = 0.645 s.

At all the time instants considered, in H-I and H-II configurations the velocity fields is almost stable with little fluctuations, in contrast with H-III case where, in the descending portion, velocity fluctuations are more visible. Regarding the aneurysmatic configurations, Figure 7 shows larger values of ensemble velocity SD for all the diseased scenarios at each time instant, especially during the deceleration phase of the cycle, where SD reaches values even up to about 25% of the maximum ensemble velocity. Specifically, the highest SD values are observed in the aneurysmal sac, where, due to the sudden change of morphology, the fluctuations are maximum. Among the DTAA cases, C-III seems to be the one that most favours transition to turbulence.

#### 3.2.3 Oscillatory Shear Index and von Mises Stresses

In Figure 8a we compare the ensemble OSI for all the pathological configurations with their healthy counterparts. We notice that, in all the diseased scenarios, OSI assumes significantly high values (> 0.4) largely on the lateral walls of the aneurysmal sac, in good accordance with what found by Tan et al. (2009a), with the exception of C-III and D-III. Moreover, we observe that, in P and C configurations, in correspondence of the proximal and distal aneurysm necks, there are both regions of very low (< 0.1) and very high OSI values. On the contrary, in D configurations, the proximal aneurysm neck is only characterised by high OSI. In all the diseased cases, on the superior wall of the aneurysm, in corrispondence of the dome, there is a region of great OSI, which is followed further downstream by an aera with low OSI values. The other part of the geometries not affected by the disease maintain nearly the same OSI values of the healthy scenarios: high OSI values are observed in ATA, at supraortics bifurcations and on the lesser curvature side of the arch, consistently with Numata et al. (2016).

**FIGURE 8.**
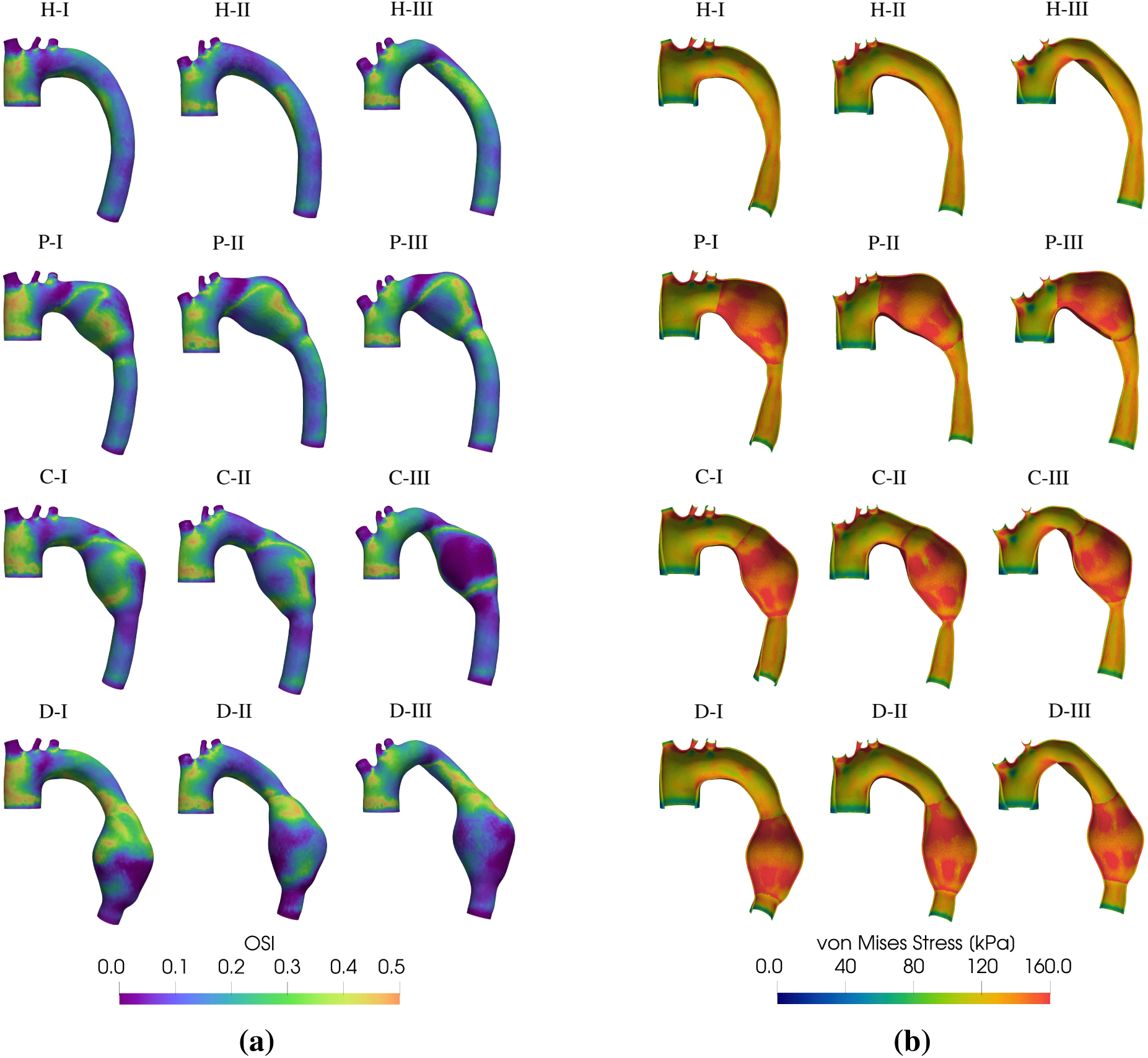
**(a)** Ensemble Oscillatory Shear Index (OSI) in all the virtual configurations. **(b)** Ensemble Von Mises Stresses (VMSs) of the aortic wall at the systolic peak (t = 0.165 s) for all the virtual configurations.

In Figure 8b we report the ensemble VMSs at the systolic peak on the aortic wall for all the virtual configurations. In the pathological cases, a clear distinction between the state of stress in the healthy portion of the geometry and in the aneurysm can be seen, due to their different Young’s moduli. In particular, in the aneurysm there is a VMSs increase up to 33% with respect to the healthy cases. The proximal and distal aneurysm necks present regions characterised by the greatest VMSs increase (almost 160 kPa), whereas the area localized at the centre of the aneurysmal sac shows relatively lower values (about 130 kPa). Also Silva et al. (2023) found that peak VMS did not occur in the larger diameter aneurysm section, but in the aneurysm neck, reporting an increase in VMS values within the Thoracic Aortic Aneurysm (TAA) from 100 kPa, in the region not afflicted by the pathology, up to 210 kPa.

### 3.3 Analysis of drag forces

In this Section, we report the ensemble drag forces results aiming to outline the changes that DTAA may cause on such hemodynamic forces, highlighting the differences between the healthy and the aneurysmatic cases, as well as the differences between each diseased scenarios, first in terms of drag forces magnitude, and then in terms of orientation.

Figure 9 shows the resultant average drag force vectors 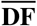 in each landing zone for all the virtual configurations, conventionally applied, for visualization purposes, at the centre point of each landing zone. From now on, in the diseased cases, we refer to those landing zones located in the non-aneurysmatic regions of TA as *standard* landing zones.

**FIGURE 9.**
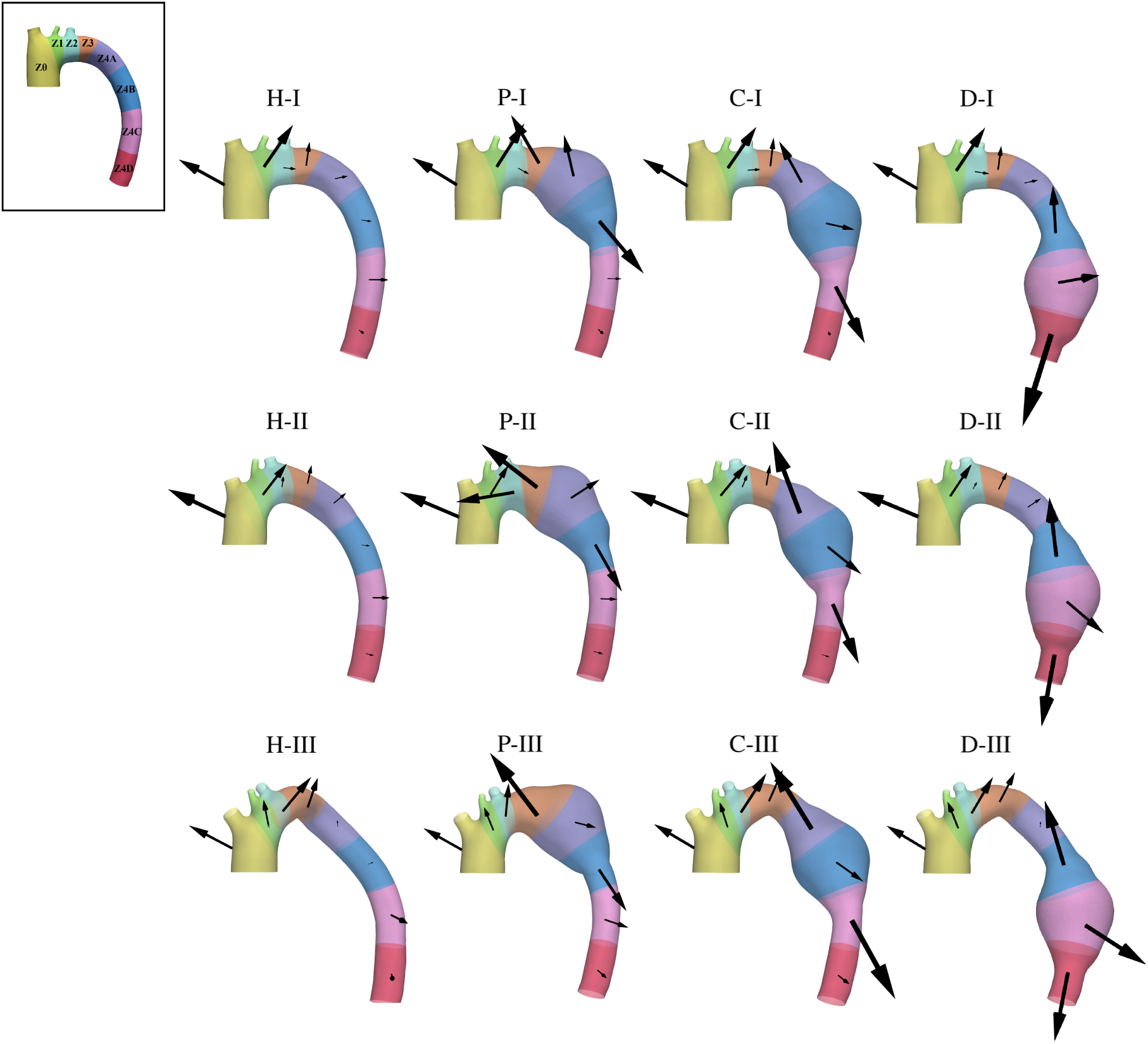
Ensemble drag forces 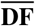 for each landing zone at the systolic peak (t = 0.165 s) for all the virtual configurations. Arrows are scaled according to 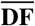 magnitude. At the top left, a representative case with landing zones is reported.

In Figure 9, we observe that, in the healthy configurations, the average drag forces with lower magnitude are located in the landing zones of the descending portion of the vessel (i.e., Z4A, Z4B, Z4C and Z4D), and this behaviour is maintained also in the diseased configurations for the standard landing zones. On the contrary, the presence of the aneurysmal sac causes a considerable 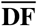 magnitude increase in the landing zones involved by the pathology. In particular, the areas with the most significant magnitude increase are the landing zones located just upstream and downstream the sac, while the increase is less notable in the landing zone ubicated in the aneurysm centre. We notice that the highest increase in 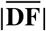, with respect to the healthy condition, is attained in D configurations. Moreover, we notice that, among all standard landing zones, Z0 is the area with the highest 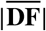 for Type II and III configurations, while it is Z1 for Type I cases. Considering the standard landing zones located at the arch (i.e., Z1, Z2 and Z3), it can be noticed that, among the three types of arch, Type III reports the lowest values of 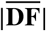 in Z1 and Z3, whereas the highest values in Z2 (except for P-II and D-III).

In Figure 9, in general we observe that the presence of the aneurysm causes the average drag forces orientation to shift gradually towards the vessel centerline direction, especially in the landing zones ubicated at the proximal and distal aneurysm necks. Regardless of the aneurysm ubication, 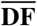 is oriented along the cranial and caudal directions (i.e., upwards and downwards) in the landing zones located just upstream and downstream the aneurysmal sac, respectively. In the aneurysm centre, in P-I scenario, 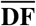 is oriented along the cranial direction, in D-I and P-II scenarios it is mainly directed sideways pointing upward, while in the other configurations it shifts more downwards. Moreover, cosidering the healthy configurations, in the landing zones situated in the descending part of the vessel, the average drag forces are mainly oriented towards the aortic wall (except for Z4A in H-III), and this behaviour is maintained also in standard landing zones in the aneurysmatic scenarios. In addition, we observe that, in all the healthy and diseased configurations, 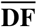 in Z0 maintains approximately the same orientation: sideways and directed towards the heart. Finally, regarding the standard landing zones ubicated at the arch, the aortic curvature increase causes the orientation of 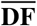 in Z1 and Z2 to shift gradually upwards, while the opposite happens in Z3.

Analysing more in detail how drag forces are oriented, in Table 5 we report the values of *DF*_∥_ and *DF*_⊥_, expressed in N/ m^2^, for each landing zone in all the virtual configurations. Regarding the healthy cases, we first observe that the greatest drag forces are located in the proximal landing zones with comparable values of *DF*_∥_ and *DF*_⊥_. Instead, the drag forces components seem to be elevated in presence of the aneurysm, as highlighted by the grey cells identifying the aneurysmatic landing zones. Specifically, in such regions, landing zones corresponding to the proximal and distal anuerysm necks are always characterised by a *DF*_∥_ larger than *DF*_⊥_ (with the exception of P-III). Regarding the standard landing zones ubicated at the aortic arch, we observe that the increase in arch curvature increases the number of zones in which *DF*_∥_ is predominant over *DF*_⊥_. In addition, we observe that C-III and D-III exhibit higher values of *DF*_∥_ in almost all the landing zones. In C-II and D-II, in Z2 and Z3, *DF*_∥_ is lesser than *DF*_⊥_, while the opposite happens in Z1. In C-I and D-I, *DF*_∥_ results to be higher than *DF*_⊥_ in Z1 and Z2, while it is lower in Z3. In P configurations, *DF*_∥_ is higher than *DF*_⊥_ both in Z1 and Z2.

**TABLE 5.** *DF*_∥_ and *DF*_⊥_ (expressed in N/m^2^), and the risk factor *β* (expressed in N^2^/m^4^) computed at the systolic peak (t = 0.165 s) in each landing zone for all the virtual configurations. Grey cells indicate the landing zones directly affected by the pathology. Red numbers indicate the maximum value of *β* for each of the nine diseased configurations. Green cells identify the closest proximal and distal landing zones that we suggest for the TEVAR stent-graft anchoring.

| Arch Type | LZ | H |  | P |  |  | C |  |  | D |  |  |
| --- | --- | --- | --- | --- | --- | --- | --- | --- | --- | --- | --- | --- |
| | | $DF_{\parallel}$ | $DF_{\perp}$ | $DF_{\parallel}$ | $DF_{\perp}$ | $\beta$ | $DF_{\parallel}$ | $DF_{\perp}$ | $\beta$ | $DF_{\parallel}$ | $DF_{\perp}$ | $\beta$ |
| I | Z0 | 1849.1 | 1918.8 | 1983.0 | 1824.7 | 4.2 | 1905.5 | 1884.5 | 3.7 | 1899.4 | 1831.3 | 3.7 |
|  | Z1 | 3893.8 | 1263.7 | 3918.0 | 1384.4 | 0.3 | 3891.8 | 1262.3 | 26.6 | 3800.8 | 1389.6 | 23.6 |
|  | Z2 | 837.5 | 171.9 | 691.6 | 264.2 | 0.7 | 899.1 | 108.6 | 2.4 | 868.5 | 194.8 | 1.6 |
|  | Z3 | 74.8 | 2776.3 | 3895.9 | 2173.6 | 146.7 | 235.8 | 3190.5 | 0.0 | 162.1 | 2897.9 | 0.0 |
|  | Z4A | 680.7 | 752.3 | 2430.4 | 676.0 | 21.2 | 3597.1 | 447.1 | 84.4 | 636.0 | 805.6 | 0.3 |
|  | Z4B | 402.3 | 396.3 | 4621.8 | 510.7 | 217.8 | 693.0 | 1145.3 | 0.5 | 3734.1 | 163.6 | 202.9 |
|  | Z4C | 68.7 | 965.5 | 17.9 | 709.8 | 0.0 | 2920.1 | 578.4 | 124.9 | 348.5 | 1833.9 | 0.1 |
|  | Z4D | 115.8 | 567.1 | 144.4 | 644.0 | 0.0 | 130.4 | 530.1 | 0.0 | 5188.0 | 12.3 | 3693.5 |
| II | Z0 | 1669.8 | 3852.4 | 1637.9 | 3642.9 | 1.8 | 1568.3 | 3780.3 | 1.5 | 1632.8 | 3792.7 | 1.7 |
|  | Z1 | 2848.9 | 324.4 | 2475.2 | 420.3 | 13.9 | 2951.7 | 331.5 | 26.5 | 2776.7 | 408.1 | 19.9 |
|  | Z2 | 411.5 | 469.4 | 2888.5 | 12.6 | 334.7 | 454.5 | 518.3 | 0.2 | 278.4 | 366.0 | 0.1 |
|  | Z3 | 313.0 | 2015.1 | 6312.4 | 420.3 | 693.4 | 214.5 | 2141.8 | 0.0 | 320.6 | 1748.2 | 0.0 |
|  | Z4A | 437.2 | 1046.1 | 844.8 | 1515.0 | 0.7 | 4870.1 | 877.9 | 186.5 | 438.0 | 960.8 | 0.1 |
|  | Z4B | 165.3 | 528.6 | 3546.7 | 471.4 | 159.8 | 2072.2 | 826.3 | 24.1 | 4780.2 | 52.1 | 1176.6 |
|  | Z4C | 53.4 | 1062.1 | 53.9 | 936.0 | 0.0 | 3068.6 | 286.1 | 233.8 | 1269.3 | 920.6 | 9.2 |
|  | Z4D | 49.5 | 491.3 | 71.5 | 505.2 | 0.0 | 48.8 | 392.4 | 0.0 | 3729.6 | 179.6 | 550.5 |
| III | Z0 | 928.7 | 2729.4 | 1002.4 | 2564.0 | 0.7 | 1068.8 | 2491.2 | 0.8 | 1070.6 | 2417.8 | 0.8 |
|  | Z1 | 1592.6 | 1415.6 | 1491.7 | 1469.8 | 2.2 | 1603.9 | 1309.7 | 2.9 | 1559.9 | 1570.0 | 2.4 |
|  | Z2 | 1872.9 | 568.2 | 1185.1 | 1085.6 | 1.2 | 2195.0 | 583.6 | 10.1 | 1965.4 | 569.5 | 7.4 |
|  | Z3 | 1394.9 | 903.9 | 2667.7 | 3306.1 | 8.8 | 1333.0 | 955.0 | 2.1 | 1476.5 | 18.2 | 20.2 |
|  | Z4A | 443.2 | 175.3 | 1082.4 | 898.1 | 2.0 | 5757.0 | 125.1 | 810.4 | 383.5 | 182.2 | 0.2 |
|  | Z4B | 27.9 | 379.6 | 3227.3 | 569.5 | 266.8 | 1362.1 | 656.8 | 18.7 | 4207.3 | 697.5 | 534.1 |
|  | Z4C | 66.3 | 1455.4 | 60.1 | 1362.4 | 0.0 | 3494.8 | 958.3 | 169.3 | 1151.2 | 1797.6 | 4.4 |
|  | Z4D | 108.4 | 1016.6 | 92.2 | 856.4 | 0.0 | 90.7 | 900.7 | 0.0 | 3644.1 | 8.0 | 1641.3 |

In Table 5, we report also the risk factor *β* evaluated for each landing zone in all the diseased configurations. Consistently with what observed for 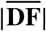 and for *DF*_∥_ and *DF*_⊥_ values, we notice that, for each configuration, the maximum value of *β* (red numbers in Table 5) is always attained in correspondence of the aneurysm (grey cells in Table 5). In particular, we observe that in six out of nine diseased cases, the highest *β* is found in the landing zone located in corrispondence of the distal aneurysm neck. In addition, the lowest *β* value, in the regions involved by the pathology, is always encountered in the landing zone ubicated at the aneurysm centre. Moreover, we notice that the highest value of *β* in the standard landing zones is always attained in the landing zones upstream the aneurysmal sac. Specifically, the standard landing zone which exhibits the highest risk factor is Z1 for C-I, D-I, C-II, D-II and P-III, probably due to the fact that *DF*_∥_ results to be much larger than *DF*_⊥_. Instead, the highest values of *β* is encountered in Z2 for P-II and C-III, and in Z3 for D-III. Even though the 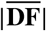 in Z0 is elevated, the risk factor associated to that landing zone remains relatively low, probably because *DF*_∥_ and *DF*_⊥_ do not substantially change with respect to the healthy counterpart, and because *DF*_⊥_ is predominant over *DF*_∥_ (except for Type I configurations in which, however, *DF*_⊥_ is comparable to *DF*_∥_).

Finally, in Table 5 we highlight in green the landing zones we suggest for the stent-graft anchoring in each diseased configuration, according to the risk factor *β*. In all the cases, we select the proximal and distal landing zones closest to the aneurysm which exhibits low *β* values.

## 4 RESULTS: PATIENT-SPECIFIC CONFIGURATIONS

In this Section we analyse the results of the simulations obtained for the three real cases reported in Section 2.1 (i.e., PT-09, PT-40 and PT-71), with the attempt of confirming the outcomes found and discussed for their corresponding virtual cases (i.e., C-III, D-I and P-II, respectively) in the previous Section. The aim is to verify whether predictions in terms of hemodynamic indices obtained in the ideal cases could be transferred to real aneurysmatic cases.

### 4.1 Hemodynamic indices

In Figure 10 we report the ensemble velocity over seven HBs for all the patient-specific configurations at two time instants: peak systole (t = 0.165 s) and late diastole (t = 0.645 s).

**FIGURE 10.**
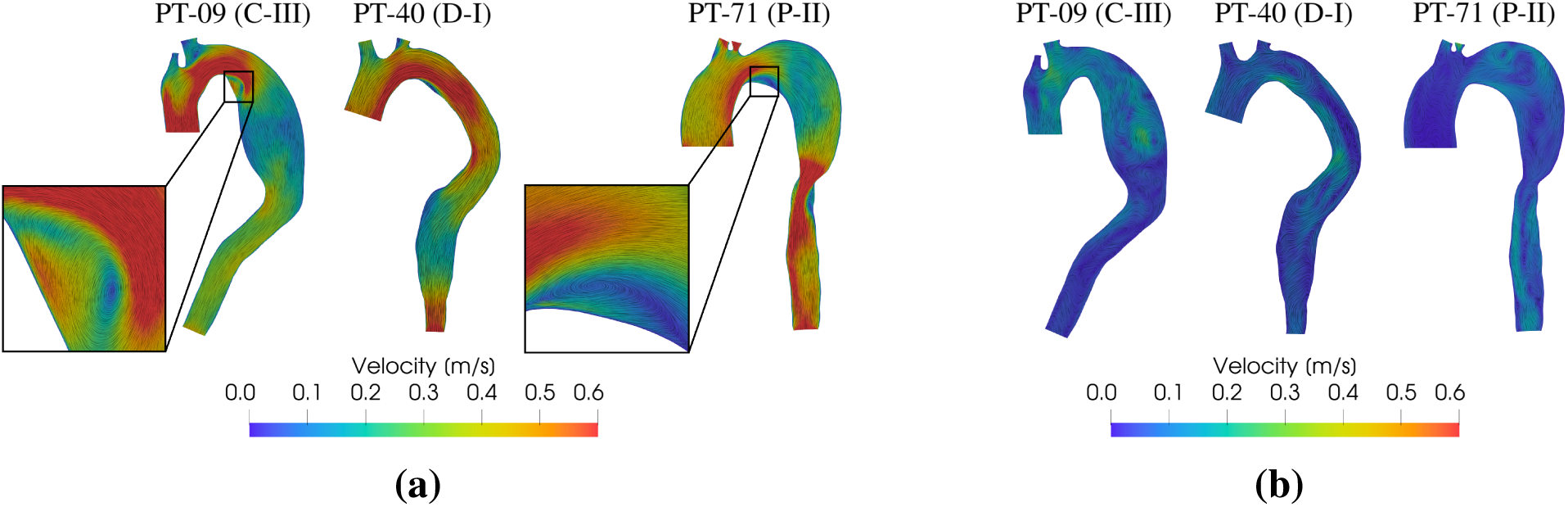
**(a)** Ensemble velocity magnitude over seven HBs at the systolic peak (t = 0.165 s) in all the patient-specific configurations. Black boxes highlight the vortex forming near the proximal aneurysm neck. **(b)** Ensemble velocity magnitude over seven HBs at late diastole (t = 0.645 s) in all the patient-specific configurations. In the brackets we report the corresponding virtual scenario.

At the systolic peak (Figure 10a), for PT-09 we can observe a velocity field very similar to the ideal C-III case (see Figure 6a): there is the presence of a high velocity jet, that impinges from the region upstream the aneurysm into the aneurysmal sac, which however disappears inside the bulging, since the latter is characterised by a considerable velocity magnitude reduction. Also for PT-40 it is possible to see great accordance with the velocity field distribution found for the virtual D-I case: a jet characterised by high velocity upstream the aneurysm and low velocity inside the aneurysmal sac. For PT-71, blood velocity significantly increases downstream the aneurysm, if compared to its corresponding virtual case P-II, probably because of the vessel shrinkage at the distal aneurysm neck. Moreover, the numerical simulations in the real scenarios are able to reproduce once again the vortical structure near the proximal aneurysm neck (black boxes in Figure 10a), which was encountered also in the ideal configurations.

In the diastolic phase (Figure 10b), in all the configurations the streamlines show chaotic behaviour, demonstrating the presence of vortices inside the aneurysmal sac. This is in accordance with what was reported for the ideal corresponding cases (see Figure 6b).

Figure 11 shows the SD of the ensemble velocity magnitude over seven HBs for all the patient-specific configurations at two different time instants: early systole (t = 0.09 s) and late diastole (t = 0.645 s). As in the virtual cases, we notice that SD is particularly high within the aneurysm, thus demostrating great velocity fluctuations and so a likely transition to turbulence. In particular, in PT-09, SD reaches values of about 28% of the maximum ensemble velocity, in good agreement with its ideal corresponding case, in which SD reached values of about 35% of the maximum ensemble velocity. Instead, PT-40 shows lower SD values if compared with the other two patient-specific scenarios, in accordance with the study in the ideal cases that showed smaller values of SD for Type I aortic arch (see Figure 7). PT-71 is characterised by high values of SD also downstream the aneurysmal sac, probably caused by the narrowing of the vessel at the distal aneurysm neck.

**FIGURE 11.**
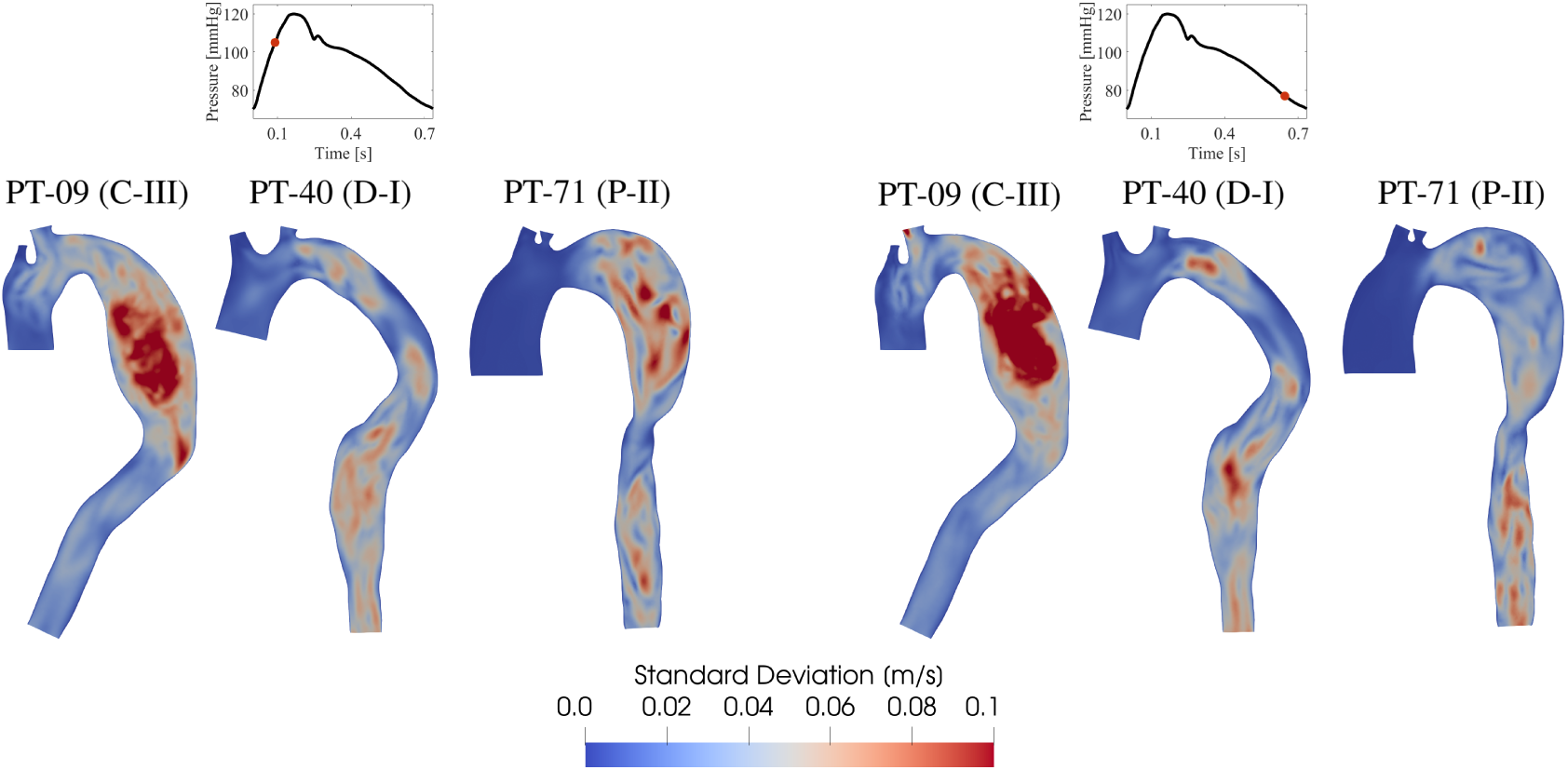
Standard Deviation of the ensemble velocity magnitude over seven HBs for all the real configurations at two time instants: (left) early systole (t = 0.09 s) and (right) late diastole (t = 0.645 s). In the brackets we report the corresponding virtual scenario.

### 4.2 Drag forces magnitude and orientation

Figure 12 shows the ensemble drag forces 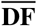 magnitude and orientation in each landing zone for all the patient-specific configurations. As it happens for the corresponding virtual cases (i.e., C-III, D-I and P-II, see Figure 9 and Table 5), the landing zones ubicated in corrispondence of the proximal and distal aneurysm necks show the greatest 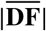. Moreover, we observe that the proximal standard landing zone which exhibits the highest 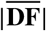 is Z0 for PT-71 (consistently with P-II), while it is Z1 for PT-09 and PT-40 (consistently with D-I). In addition, in accordance with the ideal cases, we observe that in Z1 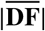 is maximum for PT-40 and minimum for PT-71, whereas in Z2 the maximum is attained for PT-09. In addition, in the landing zones ubicated in correspondence of the aneurysm necks, 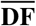 is mainly oriented towards the vessel centerline direction, consistently with the corresponding virtual scenarios. In all the patient-specific configurations, 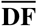 in Z0 is mainly oriented towards the heart, coerently with the virtual cases.

**FIGURE 12.**
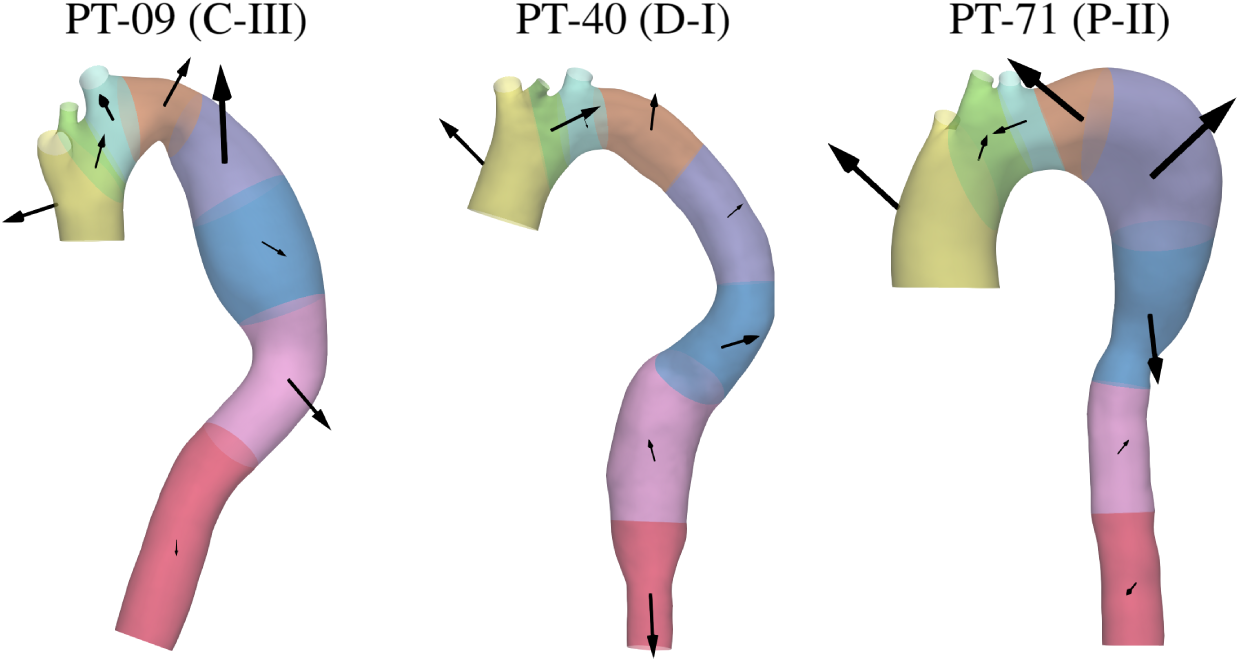
Ensemble drag forces 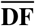 of each landing zone at the systolic peak (t = 0.165 s) for all the patient-specific configurations. Arrows are scaled according to 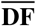 magnitude. For each patient the corresponding virtual configuration is reported in the brackets.

Regarding PT-09, 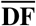 in Z4A and 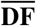 in Z4C points upwards and downwards, respectively, in good agreement with C-III. Also the orientation of 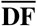 in Z4B, which is localized in the aneurysm centre, is similar with the ideal corresponding configuration. Regarding the proximal standard landing zones, 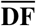 in Z1, Z2 and Z3 is mainly directed in the cranial direction, in good agreement with C-III.

In PT-40 we notice that the orientation of 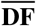 in Z4D is almost the same as in D-I, whereas this does not hold for the direction in Z4B and in Z4C, probably because of the more pronounced curvature that characterises the real DTA. However, 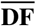 direction in the proximal standard landing zones is consistent with D-I.

For PT-71, we observe the same 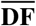 orientation of P-II in all the three landing zones directly affected by the pathology (i.e., Z3, Z4A and Z4B). In Z2, 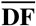 results to be mainly directed towards the heart, while in Z1 it is mainly oriented along the cranial direction, as it happens in its corresponding ideal case.

In Table 6, we report the values of *DF*_∥_ and *DF*_⊥_ in each landing zone for all the patient-specific configurations. As in the virtual scenarios (see Table 5), in the landing zones located in correspondence of the aneurysm necks *DF*_∥_ is greater than *DF*_⊥_ in all the three cases, except for Z4C in PT-09, probably because of the slight curvature of its DTA. In the landing zone ubicated at the aneurysm centre, *DF*_∥_ results to be larger than *DF*_⊥_ in PT-40 and PT-09 (as in C-III), in contrast to PT-71, which is however consistent with what observed for P-II. Regarding the standard landing zones located at the aortic arch, in Z0 *DF*_⊥_ results to be higher than *DF*_∥_ in PT-09 (as in C-III) and PT-40 but not in PT-71. In Z1 *DF*_∥_ is predominant over *DF*_⊥_ in all the three cases (as in the virtual scenarios), and in Z2 *DF*_∥_ exceeds *DF*_⊥_ in PT-71 (as in P-II), while the opposite happens in PT-09 and PT-40. In Table 6 we report also the risk factor *β* in each landing zone for all the three real configurations. Consistently with the ideal cases, the highest risk factor values (red numbers in Table 6) are always found in corrispondence of the aneurysmal sac (grey cells in Table 6). Moreover, the standard landing zone with the highest value of *β* results to be Z1 for PT-09 and PT-40 and Z2 for PT-71, consistently with their virtual counterparts (except for PT-09).

**TABLE 6.** *DF*_∥_ and *DF*_⊥_ (expressed in N/m^2^), and the risk factor *β* (expressed in N^2^/m^4^) computed at the systolic peak (t = 0.165 s) in each landing zone for all the patient-specific configurations. Grey cells indicate the landing zones directly affected by the pathology. Red numbers indicate the maximum value of *β* for each of the three diseased configurations. Green cells identify the closest proximal and distal landing zones that we suggest for the TEVAR stent-graft anchoring. In the brackets we report the corresponding virtual scenario.

| LZ | PT-09 (C-III) |  |  | PT-40 (D-I) |  |  | PT-71 (P-II) |  |  |
| --- | --- | --- | --- | --- | --- | --- | --- | --- | --- |
| | $DF_{\parallel}$ | $DF_{\perp}$ | $\beta$ | $DF_{\parallel}$ | $DF_{\perp}$ | $\beta$ | $DF_{\parallel}$ | $DF_{\perp}$ | $\beta$ |
| Z0 | 705.2 | 2221.9 | 0.2 | 608.0 | 2037.6 | 0.1 | 2002.6 | 1387.0 | 5.3 |
| Z1 | 1841.9 | 1099.5 | 4.7 | 2960.0 | 100.2 | 41.5 | 1011.6 | 631.7 | 0.8 |
| Z2 | 683.7 | 2141.6 | 0.2 | 140.4 | 399.3 | 0.0 | 2213.9 | 108.6 | 51.3 |
| Z3 | 742.6 | 2078.4 | 0.2 | 622.1 | 1188.5 | 0.8 | 3596.3 | 1330.4 | 72.1 |
| Z4A | 3615.6 | 739.6 | 82.6 | 23.3 | 739.5 | 0.0 | 823.7 | 2217.6 | 0.6 |
| Z4B | 827.4 | 206.8 | 7.5 | 1588.6 | 526.2 | 8.7 | 2286.4 | 280.5 | 55.5 |
| Z4C | 481.9 | 2065.9 | 0.3 | 586.7 | 243.8 | 1.6 | 336.3 | 231.6 | 0.3 |
| Z4D | 289.7 | 50.3 | 0.3 | 2162.5 | 13.2 | 258.6 | 243.2 | 160.6 | 0.2 |

In Table 6, as a very preliminary result, we highlight in green the landing zones we suggest for the stent-graft anchoring in all the three patient-specific configurations. As done for the virtual cases, we choose as possible landing zones those closest to the aneurysmal sac and with the lowest value of *β*. Notice the excellent agreement with the study of the ideal cases.

## 5 DISCUSSION

### 5.1 On the novelties of the study

This study presents a detailed description of the hemodynamics and wall state of stress of Descending Thoracic Aortic Aneurysms (DTAAs) in virtual and patient-specific scenarios, also provinding valuable insights on the forces exerted by the blood flow on the aortic wall. To do this, we performed a 3D Fluid-Structure Interaction (FSI) analysis with a turbulence model, to capture transition to turbulence highly characterising thoracic aortic blood flow, and distinguishing the elastic properties of the healthy and aneurysmatic portion of the vessel.

To the best of our knowledge, this is the first attempt where such a sophisticated model has been used to obtain relevant outcomes in terms of forces distribution in DTAAs. Moreover, this is the first time where an hemodynamic analysis has been performed for several DTAA configurations with different aneurysm locations and aortic arch types.

### 5.2 On the physiological and pathophysiological relevance of the study

The presence of the aneurysm causes an increase of peak pressure both within the aneurysmal sac and in the downstream portion of Descending Thoracic Aorta (DTA), while it provokes a sudden change in pressure values at the proximal and distal necks of the bulging (see Figure 5a). These results are in good agreement with the computational studies conducted by Etli et al. (2021) and by Dadras et al. (2023). The former reported a pressure increase from a normal subject to two TAA cases during the systolic peak, while the latter showed a homogeneously higher pressure within the aneurysm, with a sudden change in pressure values at the distal neck of the sac. We argue that the peak pressure increase could be due to the higher stiffness imposed to the aneurysmal wall, consistently with what found by Campobasso et al. (2018). Considering the increase of peak pressure inside the aneurysmal sac, we can presume that the presence of DTAA may enhance the load on the aortic wall, and thus, potentially increase the risk of aneurysm enlargement and rupture (Choke et al., 2005).

Thanks to our model, which allows us to account for different Young’s moduli for the healthy and aneurysmatic portion of the vessel, we obtain that the middle cross-sectional area variation within one heartbeat (HB) is halved when passing from the healthy to the diseased Thoracic Aorta (TA) (see Figure 4b). In addition, given the higher stiffness of the aneurysmal sac, the aneurysmatic wall is subjected to greater internal stresses, in accordance with the Von Mises Stresses (VMSs) distribution (see Figure 8b), increasing the likelihood of rupture. In particular, our numerical results show that the most stressed regions are the proximal and distal aneurysm necks, in which therefore we speculate that the rupture is more likely to occur, whereas the area at the centre of the aneurysmal sac shows lower VMS values. These results are in accordance with what reported by Silva et al. (2023), who found that peak VMS did not occur in the larger diameter aneurysm section, but at the aneurysm necks.

At the systolic peak, within the aneurysmal sac, blood flow is characterised by disordered streamlines, demonstrating the presence of tiny vortices on the lesser curvature side near the proximal aneurysm neck (see Figure 6a), which might be regions of blood recirculation. These recirculation regions favour the formation of blood stagnation (Ong et al., 2019) and increase in size with the degree of aortic arch curvature (i.e., from arch Type I to III), resulting more evident in P-III and C-III. Besides, C-III scenario presents also another vortical structure within the aneurysmal sac, further enhancing the blood recirculation. The latter is strictly related to aneurysm rupture (Numata et al., 2016) and Intraluminal Thrombus (ILT) formation (Ong et al., 2018), confirming that sharp curvature (i.e., Type III), at the inner side wall of the aortic arch, causes flow separation and the rolling up of the shear layer to form conditions facilitating ILT formation and thus aneurysm rupture (Numata et al., 2016; Ong et al., 2018). In the same regard, Arzani et al. (2014) found that very low Oscillatory Shear Index (OSI) values (< 0.1) may identify regions where ILT formation is more likely to occur. Interestingly, the aneurysmal sac in C-III is characterised entirely by very low OSI values. More generarly, in P and in C configurations low OSI values are attained in the regions where the recirculations are observed. On the other hand, D configurations, which instead exhibit greater values of OSI in the region near the proximal aneurysm neck, show indeed recirculation regions with a much smaller size in such area.

On the other side, the OSI distribution exhibits high values in several regions within the aneurysmal sac and at the aneurysm’s necks in all the configurations (see Figure 8a), in good agreement with Numata et al. (2016). This means that all the configurations under study may be at risk in terms of atherosclerotic plaque formation (Zeng and Li, 2013). This is a serious risk as plaque can burst or rupture the inner wall intima continuing up to the adventitia (Febina et al., 2018). Specifically, we notice that for proximal aneurysms (P configurations) it seems that the regions most at risk is the aneurysmal sac, whereas for distal aneurysms (D configurations) the region most at risk reveals to be the proximal aneurysm neck or even the healthy portion of DTA upstream the aneurysm (D-III).

The distribution of the Standard Deviation (SD) of the ensemble velocity magnitude allows us to quantify and localize the velocity fluctuations among the HBs and thus regions of transition to turbulence (Vergara et al., 2017). SD reaches its highest values within the aneurysm (see Figure 7), confirming that transition to turbulence develops inside the sac, especially during blood deceleration phase. This disordered behavior of the velocity field may be caused by the sudden expansion of the flow stream, due to abrupt vessel enlargement in correspondence of the aneurysmal sac (Tan et al., 2009a). Turbulent flow could generate higher stresses on the aneurysm wall, if compared with laminar flow, causing an increasing rate of wall dilation, thus enhancing turbulence itself, establishing a mechanism for aneurysm dilation (Khanafer et al., 2007). We also notice that, as the aortic arch curvature increases, the flow regime becomes more unstable and tends more towards a regime of transition to turbulence. Indeed, even in the healthy case H-III it is possible to identify a region with high turbulence in the descending portion of the vessel. Probably, in Type III configurations, the steep aortic arch curvature provokes an high velocity jet in the areas upstream the aneurysmal sac, which promotes transition to turbulence (Tan et al., 2009a).

### 5.3 Towards preliminary indications for TEVAR procedure

Drag forces were intensively analysed to assess the stress state of the aorta after TEVAR implantation Fung et al. (2008); Figueroa and Zarins (2011); Krsmanovic et al. (2014). Here, we want to investigate such quantites to have an idea of the state of stress of aneurysmatic cases before the insertion of the stent-graft, with the hope that this analysis could be of support for TEVAR planning and outcomes prediction, provided that future studies will be addressed to establish such relationship. In this study, we have evaluated the drag forces magnitude and orientation in each landing zone intended by the MALAN classification (Marrocco-Trischitta et al., 2017).

The comparison between aneurysmal and healthy TA showed no differences in terms of drag forces magnitude and orientation in the landing zones not directly involved by the pathology (i.e., standard landing zones). Instead, the presence of the aneurysmal sac causes an increase in drag forces magnitude and a gradual shift of their orientation towards the vessel centerline in the landing zones ubicated at the proximal and distal aneurysm necks. In particular, in such zones, drag forces magnitude results to be from 3 up to 6 times higher than those in the adjacent zones, consistently with the pressure distribution and with what reported by Marrocco-Trischitta et al. (2019). Moreover, considering the standard landing zones, the drag forces with the greatest magnitude are localized in the proximal landing zones involving the aortic arch (i.e., Z1, Z2 and Z3), in good accordance with what found by Marrocco-Trischitta et al. (2018). The landing zones ubicated at the aortic arch exhibit larger values of the resultant drag forces, because of the restricted surface area and the higher pressure values that characterise the former zones. The resultant drag forces orientation largely depends on the aortic anatomic features (Figueroa et al., 2009; Kandail et al., 2014; Marrocco-Trischitta et al., 2018). Indeed, changes in the shape of a certain landing zone cause changes in the direction of the surface normals, consequently leading to a shift of the resultant drag force orientation. For instance, if the landing zone presents some curvatures (such as the aneurysm neck), there is a deflection of the resultant drag forces orientation.

In what follows, we go deeper in our pre-operative analysis to understand whether and how it could provide some preliminary results in relation to the identification of the best landing zones for the stent-graft sealing. This could be in future very useful for clinicians in order to have information on how to perform the anchorage before TEVAR procedure. According to Domanin et al. (2021), during the planning phase for TEVAR procedure, it may be useful to estimate in advance the hemodynamic drag forces, due to the blood flow, acting on the aortic wall. Indeed, a notably challenging hemodynamics, typical of certain regions of the aneurysmatic DTA, may cause the generation of particularly hostile drag forces, which potentially lead to an insufficient proximal or distal sealing likely causing the stent-graft migration (Figueroa et al., 2009).

In order to take into account all the threatening conditions that may lead to endograft migration (see Section 2.5.2), we propose a risk factor *β*: a single index able to summarizes all the dangerous conditions in order to classify each landing zone according to its risk of migration. The greatest value of *β* is always attained in the landing zones located in corrispondence of the proximal and distal aneurysm necks. This behaviour could be explained by two mechanisms: first, because in such regions the drag forces component parallel to the vessel centerline is always greater than the normal one; second, because of the noticeable variation of the drag forces with respect to the healthy condition. For these reasons, this landing zones should be avoided for the stent-graft sealing. Moreover, the highest value of *β* in the standard landing zones is always attained in the landing zones upstream the aneurysmal sac. Indeed, in these regions, it is often the case that the parallel component results to be larger than the perpendicular one, even though there is no substantial change between the diseased and the corresponding healthy cases. In particular, for Type III configurations, the parallel component is always predominant over the perpendicular one in all the three landing zones ubicated at the aortic arch, even though its values are not always the highest if compared with Type I and II cases. According to *β*, the risk of endograft migration in Z1 decreases when passing from Type I to Type III aortic arch, whereas it is maximum in Z2 and Z3 for Type III configurations, except for P scenarios where the maximum and minimum risk is reached in P-II and P-I, respectively. Our results are also in agreement with the less frequency of TEVAR complications observed for proximal sealing performed in Z0 (Melissano et al., 2007). However, it is important to recall that an anchorage at that level would require the bypass of all three supraortic vessels, making the surgery more challenging (Milewski et al., 2010).

Considering the aforementioned observations, regarding the proximal stent-graft sealing (see green cells in Table 5), we notice that:

For D configurations, regardless the arch type, it would be better to choose Z4A as proximal landing zone, as it is not located at the aortic arch and it has always a perpendicular component larger than the parallel one. However, we notice that, especially for D-III, Z4A features large values of OSI, i.e., high risk of plaque formation; for this reason, the effective choice of this landing zone for the stent-graft anchoring should account this risk;

- For P-I, P-III and C-II configurations, Z2 seems to be the most suitable proximal landing zone, since it shows a slight difference between the parallel and the perpendicular component, both with reduced magnitude, if compared with the neighbouring landing zones;
- For P-II configuration, Z0 could be the best choice for the proximal sealing, since it has the lowest *β* value, while Z1 and Z2 are characterised by a too high parallel component in relation to the perpendicular one;
- For C-I and C-III configurations, the better choice could be Z3 because, in the former case, it has a negligible value of *β*, while, in the latter case, because it exhibits the lowest parallel and perpendicular components combination with respect to the nearby landing zones.

### 5.4 On the validity of the analysis on patient-specific cases

In this study, we have applied our computational model also to three patient-specific configurations, showing that the model and the numerical results obtained are also valid for real cases. In particular, it is possible to observe that the main features found for the ideal cases, i.e., the high velocity jet in the upstream region of the aneurysmal sac, the recirculation area near the proximal aneurysm neck at the systolic peak, as well as the relatively high velocity vortex at the late diastole, hold true for patient-specific scenario. Moreover, all the three patient-specific configurations are characterised by high SD values at different time instants of the HB, in good accordance with their corresponding virtual cases. Finally, the analysis in the real cases highlights an excellent agreement regarding the distribution of forces along TA, with respect to the ideal cases analysis. Ideed, the main results on drag forces, underlined for the virtual cases, are still valid for the real patient-specific configurations.

Specifically, we found that, also for the real cases, Type III aortic arch (PT-09) is the most prone to transition to turbulence, confirming what observed in the ideal cases, particularly in C-III (see Figure 11). Moreover, the results of the index *β* (see Table 6) confirm that, for all the three cases, maximum values are attained for the aneurysmatic landing zones (grey cells in Table 6) and the suggested landing zones for the stent-graft anchoring, characterized by low values of *β* (green cells in Table 6), are in accordance with the results of the ideal cases (see Table 5).

These findings have a great clinical relevance. Indeed, once the nine diseased cases have been simulated and analysed, it is possible to make a general *a priori* assessment of the hemodynamics and internal wall stresses of any real patient-specific case, on the basis of the virtual cases analysis.

### 5.5. Major outcomes, limitations and future developments

In what follows, we try to underline the major outcomes emerged by our analysis:

The aneurysmal sac is characterised by higher peak pressure values and von Mises stresses with respect to the healthy condition, provoking large loads in the internal wall thus inducing a risk of aneurysm rupture. Moreover, the aneurysm shows very low values of blood velocity and several blood recirculation regions, associated with low values of OSI, increasing the risk of ILT formation. On the other hand, other regions of the aneurysm are associated with large OSI values, increasing the risk of atherosclerotic plaque formation. This may be relevant in the proximal aneurysm neck and in the healthy portion of DTA upstream the aneurysmal sac for D configurations;

- Configurations with a more steep aortic arch (i.e., Type III) present a more chaotic hemodynamics, with disordered streamlines and higher likelihood of transition to turbulence. The latter may be an index of increased rupture risk for Type III. However, pressure results suggest that Type I and II could be the most critical;
- The presence of the aneurysmal sac causes the increase of ensemble drag forces magnitude, with respect to the healthy condition, and the shift of their orientation towards the vessel centerline. The greatest drag forces magnitude increase, with respect to the healthy condition, are attained in configurations with a distal aneurysm. Regarding the regions not directly affected by the pathology, the highest drag forces magnitude are encountered in the landing zones upstream the aneurysmal sac. In such regions, it is common for the drag forces parallel component to exceed the normal one;
- The risk of endograft migration at the level of the proximal sealing decreases as the aortic arch curvature increases (i.e., from Type I to Type III) in Z1, while it is maximum in Z2 and Z3 for Type III configurations;
- The numerical results found for the real patient-specific cases are in good agreement with the results of the virtual cases.

This study presents some limitations which are summarized below:

- The pressure-wave employed as inlet boundary condition for the real cases is taken from literature. A future development to assess the validity of the ideal analysis for real cases could be the inclusion for the latter of a patient-specific inlet condition derived form clinical measurements (e.g., blood flow rate or pressure over time);
- The model used for the mathematical representation of the material of the aortic wall is the linear elasticity. However, more sophisticated and realistic material models could be used, such as hyperelasticity or a fiber-based model. In addition, in this work, blood is considered as a Newtonian fluid, even though it is known that blood is a non-Newtonian, shear-thinning liquid (Gasser et al., 2024). For this reason, in the future, to better represent the hemodynamics at low shear rate (typical of the aneurysmal sac), one can opt for non-Newtonian models (Berguer et al., 2006);
- The virtual cases are willingly designed with fusiform aneurysms and with low DTA tortuosity, in order to evaluate the influence of the type of arch and ubication of the aneurysm only. In the future we plan to analyse other anatomical complexities, such as DTA tortuosity or saccular aneurysms;
- Regarding the real cases, the lack of follow-up data does not make it possible to validate our prediction regarding the possible TEVAR procedure outcomes. Thus, for the future, it will be needed to analyse cases whose evolution, in terms of possible TEVAR migration, is known;
- The risk factor *β* is designed to consider only the threatening conditions that could lead to stent-graft migration. In the future, it will be interesting to assess the risk of endoleak formation (e.g., bird-beak configuration), by designing a new appropriate index or by adapting *β*;
- The assessment of the risk of stent-graft migration is made only on the basis of the hemodynamic drag forces. As future development, it could be of interest to consider also structural parameters (e.g., von Mises Stress).

In conclusion, we believe that the analysis presented in this paper could add a new piece to the mosaic of the research to better understand the pathophysiology in DTAAs and to provide preliminary information on TEVAR implant.

## ACKNOWLEDGMENTS

FD, CV, LC are members of the INdAM group GNCS “Gruppo Nazionale per il Calcolo Scientifico” (National Group for Scientific Computing). CV has been partially supported by: i) the Italian Ministry of University and Research (MIUR) within the PRIN (Research projects of relevant national interest) MIUR PRIN22-PNRR n. P20223KSS2, “Machine learning for fluid structure interaction in cardiovascular problems: efficient solutions, model reduction, inverse problems”; ii) the Italian Ministry of Health within the PNC PROGETTO HUB LIFE SCIENCE - DIAGNOSTICA AVANZATA (HLS-DA) “INNOVA”, PNCE3-2022-23683266, CUP: D43C22004930001, within the “Piano Nazionale Complementare Ecosistema Innovativo della Salute”, Codice univoco investimento: PNCE3-2022-23683266; iii) Italian Ministry of Health within the project “CAL.HUB.RIA” - CALABRIA HUB PER RICERCA INNOVATIVA ED AVANZATA, Code: T4-AN-09, CUP: F63C22000530001.

## LIST OF SYMBOLS

| Symbol | Description |
| --- | --- |
| ALE | Arbitrary Lagrangian-Eulerian |
| ATA | Ascending Thoracic Aorta |
| BC | Boundary Condition |
| BCA | Brachiocephalic Artery |
| BDF1 | Backward Differentiation Formula of first-order |
| BDF2 | Backward Differentiation Formula of second-order |
| BL | Boundary Layer |
| CFD | Computational Fluid Dynamics |
| CTA | Computed Tomography Angiography |
| DTA | Descending Thoracic Aorta |
| DTAA | Descending Thoracic Aortic Aneurysm |
| FSI | Fluid-Structure Interaction |
| HB | Heartbeat |
| ILT | Intraluminal Thrombus |
| LCCA | Left Common Carotid Artery |
| LES | Large-Eddy Simulation |
| LSA | Left Subclavian Artery |
| MALAN | Modified Arch Landing Areas Nomenclature |
| OSI | Oscillatory Shear Index |
| SD | Standard Deviation |
| TA | Thoracic Aorta |
| TAA | Thoracic Aortic Aneurysm |
| TEVAR | Thoracic Endovascular Aortic Repair |
| 3-EWM | Three Elements Windkessel Model |
| VMS | Von Mises Stress |
| WSS | Wall Shear Stress |

## ADDITIONAL INFORMATION

### Ethical approval

this study has been approved by “Comitato Etico Milano Area 2”, protocol number: 3457/2023;

